# Endogenous Promotor-Driven Split Nanoluciferase Biosensor for Assessing G Protein Recruitment

**DOI:** 10.1101/2024.06.03.597093

**Authors:** Laura J. Humphrys, Carina Höring, Albert O. Gattor

## Abstract

HEK293 cells are a common immortal cell line used in biological research, and their popularity has led to different distinct lineages across the world. Commonly used for overexpression of proteins, HEK293 cells also natively express biological targets, such as G protein coupled receptors (GPCRs) and their downstream signalling partners, G proteins, although this often confounds rather than compliments research. CRISPR/Cas9 gene editing can be used to harness these native proteins and make use of their presence. Here, a cost- and time-effective, plasmid-based CRISPR/Cas9 approach is used to tag well-characterised GPCRS – the β-adrenoceptors 1 and 2 – with one part of a split Nanoluciferase and replace the G_αs_ coupling partner with the complimentarily tagged minimal G_s_ protein in HEK293T cells. Compared to untagged proteins, the CRISPR/Cas9 cells allow for better selective-ligand characterisation at the native β-adrenoceptors. Overexpressed tagged systems produce similar results to the CRISPR/Cas9 cells, however subtle changes in the characterisation of partial agonists, such as salbutamol, demonstrate the potential for utilising tagged native receptors in analysing biological effectors.

**Summary Statement:** For the first time, a split-luciferase tagged minimal Gs protein and β_1_AR is inserted under endogenous promotors in HEK293T cells using CRISPR/Cas9 gene modification, avoiding protein overexpression in the assay.

## Introduction

In G protein-coupled receptor (GPCR) research, split luciferase complementation (SLC)-based G protein biosensors detect protein-protein interactions in cells and rely on genetically encoded reporter proteins that are conjugated to the proteins of interest, such as the GPCR and G protein (Höring et al., 2020; Pottie et al., 2020; Vasudevan and Stove, 2020; Wan et al., 2018; Wouters et al., 2020). These biosensors use fused dissected luciferase fragments, such as the NlucC and NlucN combining to create NanoLuciferase (NanoLuc, Nluc). Complementation of the luciferase fragments upon protein-protein interaction restores their catalytic activity and promotes the oxidation of a substrate, followed by bioluminescence (Hattori and Ozawa, 2014). However, SLC biosensors have rarely allowed measurement of endogenous/native receptors due to the requirement of introducing extrinsic fusion proteins (White et al., 2017; White et al., 2019). Considering that overexpression of receptors could lead to a shift in the relative abundance of interacting partners, including potential partners for receptor dimerization, G proteins, or arrestins endogenously expressed in cells, receptor expression levels directly impact measurement of receptor reserve and signal amplitudes during ligand assessment in early drug development (Soave et al., 2021; Wacker et al., 2017).

In recent years, the modification of endogenous gene loci has been simplified by the discovery of clustered regularly interspaced short palindromic repeats (CRISPR) associated protein (Cas), CRISPR/Cas, systems, facilitating programmable site-specific DNA (or RNA) cleavage (Cong et al., 2013; Horvath and Barrangou, 2010; Jinek et al., 2012). Cas endonucleases form a complex with small (around 20 bp) guide RNA (gRNA), which locates the Cas enzyme to the target site in the genome (Cong et al., 2013; Jinek et al., 2012). As a protospacer adjacent motif (PAM) site is required for the Cas to cut the DNA at the target site, non-specific sites are safe from Cas cleavage. Therefore, the highly specific and efficient CRISPR/Cas system has become a widely used method for genome editing and has already been used to generate cells endogenously promoting engineered luciferases or fluorescent proteins (Kamiyama et al., 2016; Ratz et al., 2015; Rojas-Fernandez et al., 2015; Yang et al., 2013). In particular, the engineered NanoLuc is useful for insertion into cell genomes due to its small size (19 kDa) and bright luminescence (Hall et al., 2012). For example, NanoLuc and its dissected variant split-NanoLuc or ‘NanoBiT’ (Dixon et al., 2016) have already been integrated into cell genomes, allowing the detection of protein up-/downregulation, GPCR-β-arrestin interaction and ligand binding to a GPCR under endogenous promoters (Oh-hashi et al., 2017; Riching et al., 2020; Schwinn et al., 2018; White et al., 2017; White et al., 2019).

Immortalized HEK293 cells are among the most popular human cell lines since they are easy to transfect and provide an appropriate cellular matrix of membrane lipid composition and post-translational modifications (Atwood et al., 2011; Thomas and Smart, 2005; Wiseman et al., 2020). All these variables impact receptor conformation, signalling and regulation (Guixà-González et al., 2017; Jakubík and El-Fakahany, 2021; Patwardhan et al., 2021). However, not unsurprisingly, HEK293 cells also endogenously express GPCRs, which can complicate pharmacological experiments. Due to the wide distribution of HEK293 cells across the world allowing for different lineages, analyses of mRNA expression levels and pharmacological experiments have revealed different native GPCR expression profiles (Atwood et al., 2011; Thomas and Smart, 2005; van der Hagen et al., 2014). As a prime example, for β-adrenoceptors (βAR) there is still uncertainty as to whether different HEK293 lineages functionally express solely β_2_AR or also β_1_AR. Although the functional expression of β_1_AR in HEK293 cells has been detected by mRNA levels and cAMP responses (van der Hagen et al., 2014), other studies demonstrated only β_2_AR-mediated functional responses, which could not be blocked by a β_1_AR-selective antagonist (Copik et al., 2009; Galaz-Montoya et al., 2017; Lavoie et al., 2002; Schmitt and Stork, 2000). Subtype selectivity is crucial in the pharmacology of these receptors - β_2_-adrenoceptor agonists are the gold standard for the treatment of respiratory disorders, such as asthma (Barisione et al., 2010; Waldeck, 2002), whereas β_1_-adrenoceptor antagonists are commonly used in patients with cardiovascular diseases, such as heart failure (Ripley and Saseen, 2014; Ziff et al., 2020). Studies using HEK293 cells to study βARs therefore need to account for the potential presence of both subtypes The aim of this study was 1) to investigate whether recently developed SLC-based mini-G protein sensors are useful to detect receptor activation under endogenous promotor levels, and 2) to detect whether β_1_ARs are expressed and functional in our sample of HEK293T cells. For this purpose, CRISPR/Cas9 genome editing experiments were designed to generate endogenously promoted β_1,2_AR-NlucC fusion proteins or NlucN-mini-G_s_ (NlucN fused to the minimal G_αs_ protein) for functional coupling characterisation.

## Results

### Functional Characterisation of β1,2AR Ligands with Overexpressed Receptors

To validate our mini-G (mG) protein sensors with β_1,2_AR, a range of standard agonists and antagonists with different potencies and efficacies were characterised in HEK293T cells recombinantly co-expressing NlucN-mG_s_ and either β_1_AR-NlucC or β_2_AR-NlucC fusion proteins (“overexpressed”; Figure 1, Supplementary Information (SI), Table S2). Adrenaline and isoprenaline were full agonists at both the β_1_AR and β_2_AR. In contrast, noradrenaline was a full agonist for β_1_AR, but only partial at the β_2_AR with an E_max_ of 82±1% and lower potency (pEC_50_: 6.34±0.05 (β_1_AR) vs. 5.15±0.07 (β_2_AR)). Conversely, salbutamol activated β_2_AR with greater efficacy than the β_1_AR (E_max_: 82±1% and 37±1%, respectively) and had a substantially lower potency at the β_1_AR (pEC_50_: 5.4±0.02 (β_1_AR) vs. 7.15±0.09 (β_2_AR)). Using mG_s_ sensors with overexpressed receptors in HEK293T cells, ICI118,551 (ICI), xamoterol and carvedilol were antagonists at the β_2_AR, whereas this was only the case for ICI118,551 with the β_1_AR (Figure 1B and D, SI, Table S2). Instead, xamoterol and carvedilol were weak partial agonists with E_max_ values of 14±1% and 7±1%, respectively (Figure 1A, SI, Table S2).

**Figure 1.**
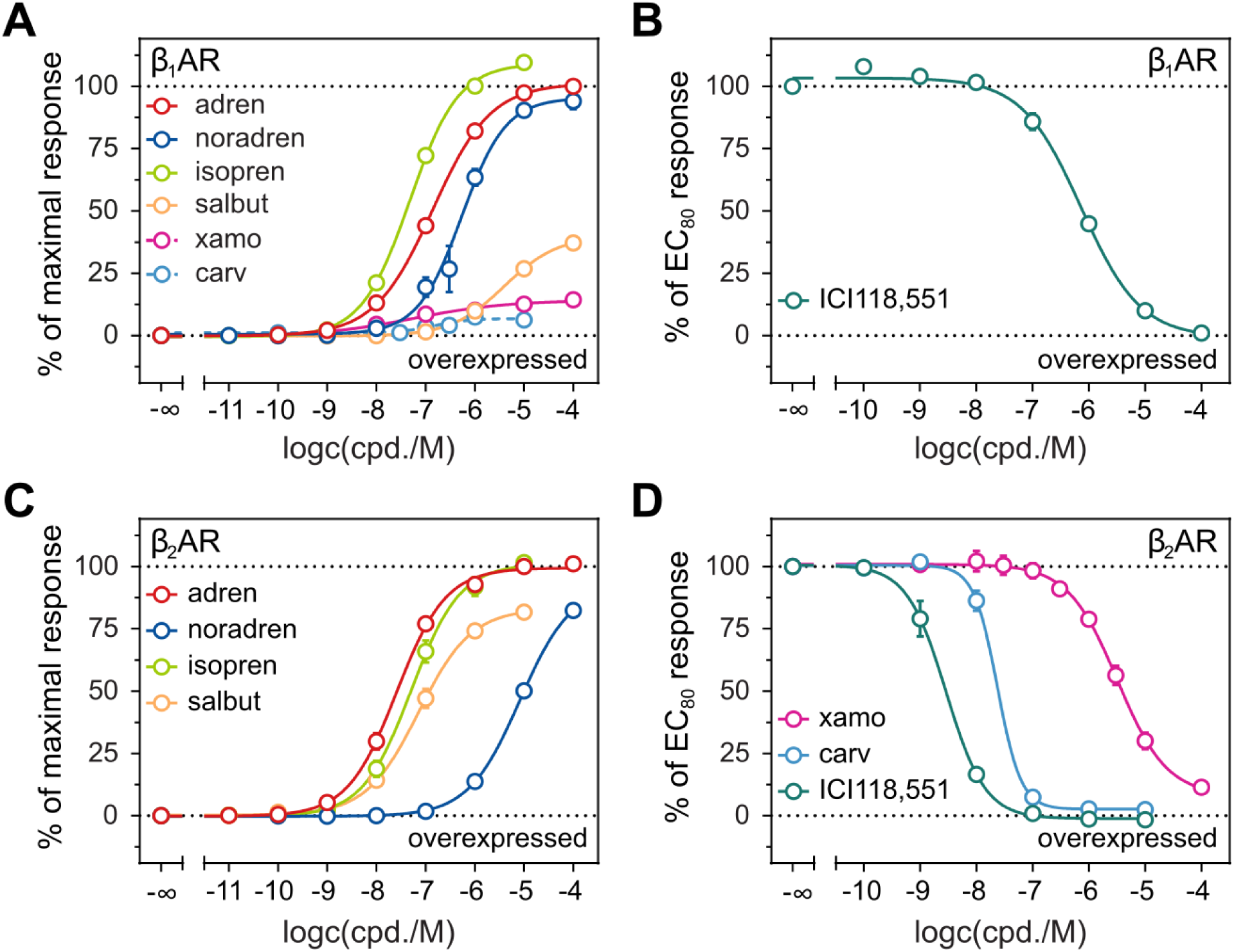
Concentration response curves of selected standard β1,2AR agonists and antagonists obtained in the mini-G protein recruitment assay. HEK293T cells recombinantly co-expressed NlucN-mGs and β1AR-NlucC (**A, B**) or β2AR-NlucC (**C, D**). Antagonists were characterised in the presence of 100 nM adrenaline (adren). Data were normalized to L-15 (0%) and to the maximal response elicited by 100 μM (β1AR) or 10 μM (β2AR) adrenaline for agonists, or 100 nM for antagonists (100%). Data represent means ± SEM from at least three independent experiments (*n* ≥ 3) performed in triplicate.

### Native Receptor Response to β1,2AR Ligands in Wildtype HEK293T Cells

Conflicting information on the expression of β-adrenoceptors in HEK293 cells exist, most likely due to the different lineages used across the world (Atwood et al., 2011; Thomas and Smart, 2005; van der Hagen et al., 2014). To assess the expression of native receptors in our HEK293T cell line, firstly, the BRET-based CAMYEN assay evaluated the cAMP production in response to β_1,2_AR ligands (Figure 2). Potencies of the agonists did not match a distinct profile of the β_1_AR or the β_2_AR. Adrenaline stimulated cAMP production with a similar potency to the β_2_AR-mG_s_ coupling (pEC_50_: CAMYEN = 7.25±0.14, β_2_AR-NluC = 7.55±0.07, β_1_AR-NlucC = 6.86±0.02), whereas noradrenaline shared a similar potency with the β_1_AR-mG_s_ coupling data (pEC_50_: CAMYEN = 6.77±0.31, β_1_AR-NlucC = 6.20±0.06, β_2_AR-NlucC = 5.12±0.06). As isoprenaline was equipotent, 100 nM isoprenaline (approximate EC_80_) was used to characterise antagonists (Figure 2B). Although ICI118,551 and carvedilol had similar potencies (pIC_50_ = 9.01±0.37 and 9.18±0.14, respectively), ICI118,551 was not able to fully antagonise the isoprenaline response up to 100 nM. When assessing the type of antagonism by using varied concentrations of isoprenaline, there was a noncompetitive effect observed for both antagonists in this assay (Figure 2C-E).

**Figure 2.**
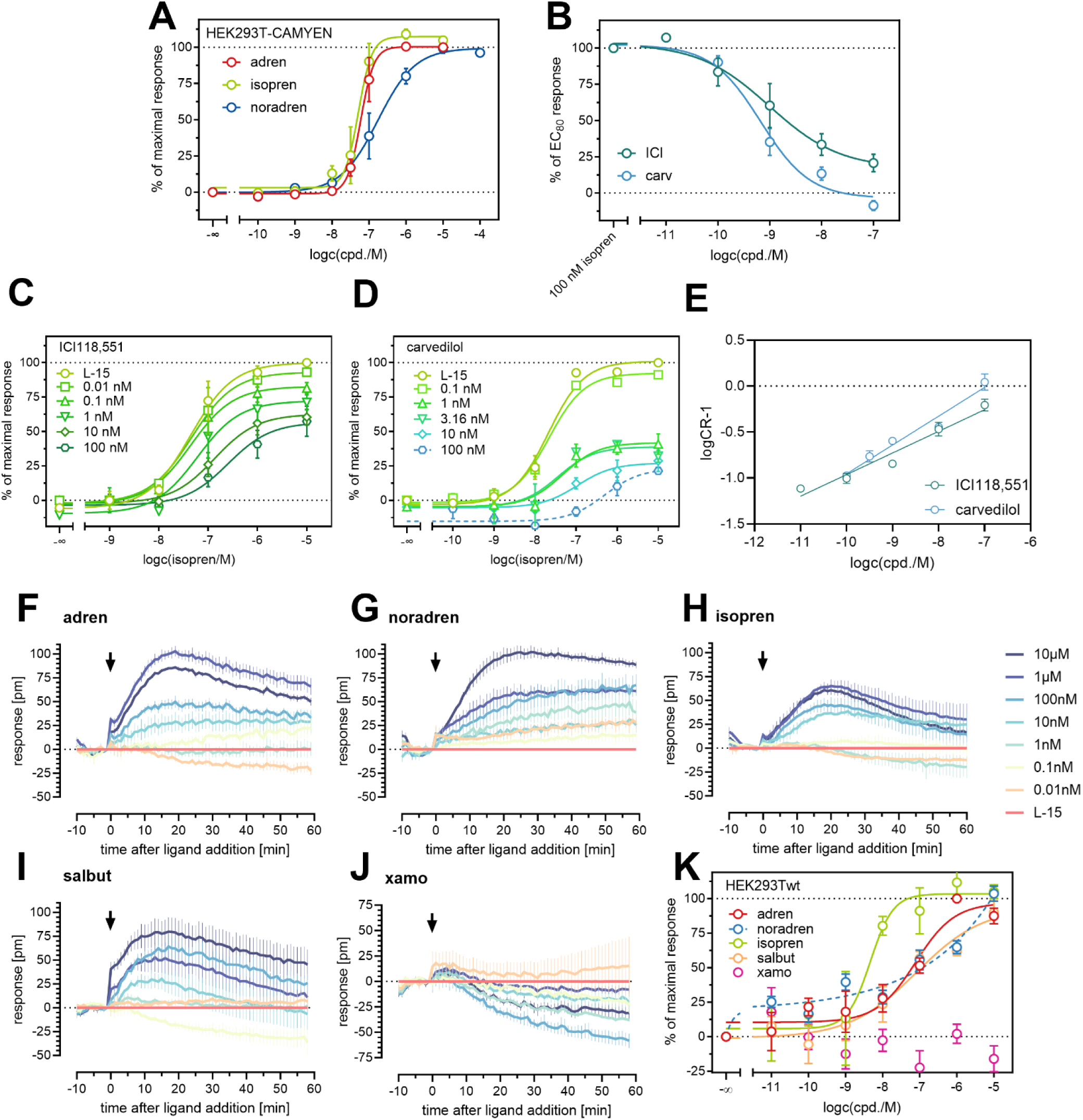
Analysis of endogenous βAR function in HEK293T cells. (**A-E**) Evaluation of cAMP responses by selected standard agonists and antagonists in wildtype HEK293T-CAMYEN cells using the BRET-based CAMYEN assay. Relative changes in [cAMP] were measured in the presence of 100 µM IBMX with 5 µM coelenterazine h. Agonist concentration response curves are shown in (A) and antagonists characterised in the presence of 100 nM isoprenaline in (B). Data were normalized to L-15 (0%) and the maximal response elicited by 10 μM adrenaline for agonists, or 100 nM isoprenaline for antagonists (100%). Antagonist responses to varied isoprenaline concentrations are shown in (C) ICI and (D) carvedilol where a three-parameter dose response curve is used. These data were used to plot the Schild linear analyses in (E), R^2^ ICI = 0.956, carv = 0.965. (**F-K**) Dynamic mass redistribution analysis of HEK293T wildtype cells with β1,2AR ligands. (E-I) represent the kinetic traces from *n* = 3 different experiments with adrenaline (E), noradrenaline (F), isoprenaline (G), salbutamol (H) and xamoterol (I). Data were corrected to cell responses to buffer only (HBSS, 0%). Note: the isoprenaline experiments were performed with different cell passages on different days to the other agonists with a reference 1 µM adrenaline response, which may impact the raw values shown. Concentration response curves in (**J**) were generated from the area under the curve of 1 h of the kinetic traces and normalised to 1 µM adrenaline (100%) and buffer (0%). Data represent means ± SEM from at least three independent experiments (*n* ≥ 3) performed in triplicate (agonists) or duplicate (antagonists).

The second functional assay was the label-free dynamic mass redistribution (DMR) assay (Schröder et al., 2011; Seibel-Ehlert et al., 2021). Here, the response to a ligand is assessed by morphological changes of the cell as downstream biochemical pathways are activated or inhibited, measured as a change in refractive index of the adherent cells by a biosensor embedded in the plate. Most agonists produced a concentration-dependent response, except for xamoterol. Noradrenaline did not produce a standard nor biphasic dose-response profile, potentially indicating the involvement of multiple receptors and/or downstream pathways. Adrenaline and isoprenaline were both comparable to the cAMP response (pEC_50_: adrenaline = 7.57±0.09, isoprenaline = 8.25±0.10). Notably, there was a potent salbutamol response (pEC_50_ = 7.06±0.41) confirming the strong presence of β_2_AR. Potency data for all experiments are summarised in the SI (Fig. S6).

Due to the discrepancies seen in the pharmacological profile of the ligands used, both assays with wildtype HEK293T cells indicated the presence of multiple βARs.

### Mini-G Protein Recruitment Assays using Endogenously Promoted β1,2AR-NlucC or NlucN-mGs

To detect β_2_AR and β_1_AR relative expressions in HEK293T cells and whether mG_s_ sensors are useful for detecting β_1_AR activation with low receptor expression (β_2_AR already validated with Nluc; Kilpatrick et al., 2019), β_1,2_AR-NlucC and NlucN-mG_s_ fusion proteins were generated by CRISPR/Cas9 genome editing of HEK293T cells (SI; Fig. S5). Responses of agonists adrenaline, noradrenaline, isoprenaline and salbutamol were probed after transient expression of NlucN-mG_s_ fusion protein in HEK293T_CRISPR_β_1,2_AR-NlucC cells. Overall, the traces of mG_s_ recruitment in response to agonists were of similar kinetics for both overexpressed and endogenously expressed β_1,2_AR (Fig. 3A, D for adrenaline). However, mG_s_ recruitment to overexpressed β_1,2_AR was faster, most likely due to the higher expression. Signal-to-background (S/B) ratios were also higher with overexpressed β_1_AR-NlucC and β_2_AR-NlucC compared to their natively promoted counterparts (Fig. 3B, E). When the sensor, NlucN-mG_s_, was endogenously promoted under the GNAS promotor, the kinetics profile changed to include a transient peak signal, more pronounced with the overexpressed β_2_AR-NlucC than β_1_AR-NlucC. The S/B ratio was further reduced with CRISPR-mG_s_ with overexpressed β_2_AR-NlucC, whereas there was no detectable difference with the native β_1_AR-NlucC S/B profiles.

**Figure 3.**
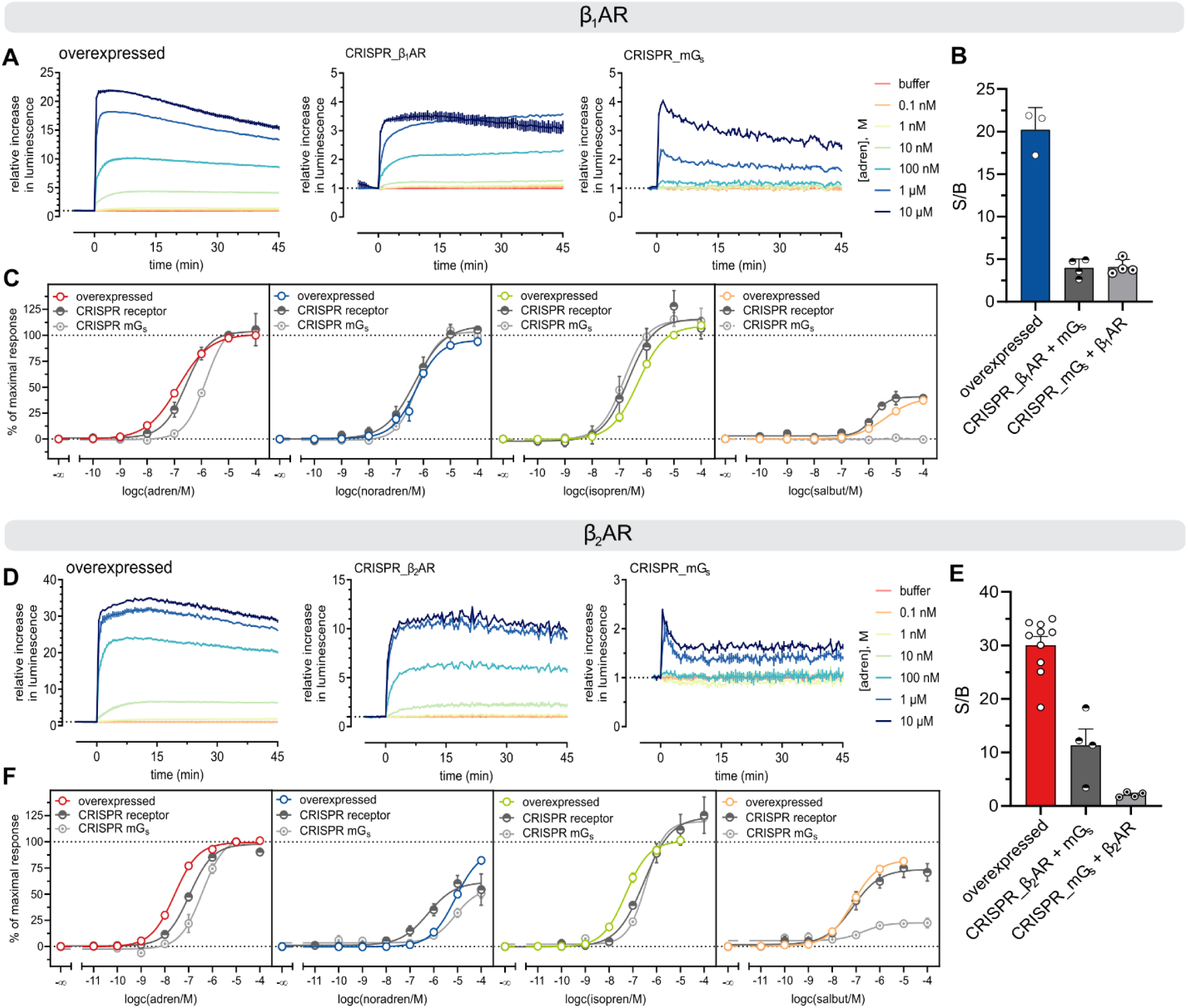
Comparison of mGs recruitment to recombinantly (overexpressed) and endogenously (CRISPR) promoted β1AR-NlucC. (**A-C**) and β2AR-NlucC (**D-F**). Representative luminescent traces of mGs recruitment (A, D) and overall signal-to-background (S/B) ratios (B, E) obtained for adrenaline are given for overexpressed and CRISPR/Cas9 modified β1AR and β2AR (96-well assays). Agonist concentration response curves were aligned for overexpressed and CRISPR/Cas9 modified β1AR (C) and β2AR (F). Mini-G protein recruitment assays were performed using HEK293T cells stably co-expressing NlucN-mGs and β1,2AR-NlucC fusion proteins, HEK293T_CRISPR_β1,2AR-NlucC that transiently expressed NlucN-mGs and HEK293T_CRISPR_NlucN-mGs that transiently expressed β1,2AR-NlucC. Data were normalized to L-15 as solvent control (0%) and to the maximal response elicited by 10 μM adrenaline, or 100 μM adrenaline for overexpressed β1AR. Data represent means ± SEM from at least three independent experiments (n ≥ 3), performed in triplicate.

Overall, agonist properties observed at the endogenous receptor level were consistent with those at overexpressed receptors. Adrenaline and isoprenaline were full agonists at both β_1,2_AR, whereas noradrenaline was a full agonist at β_1_AR but a partial agonist at β_2_AR (Fig. 3C, F). In addition, salbutamol partially activated the β_1,2_ARs. Comparing the efficacy of investigated agonists, E_max_ values were similar between overexpressed and endogenous receptors, except for noradrenaline at β_2_AR (Fig. 3; SI, Table S4). Interestingly, pEC_50_ values were increased for salbutamol at β_1_AR and noradrenaline at β_2_AR, whereas the expected pEC_50_ decrease was seen for adrenaline and isoprenaline at β_2_AR (Fig. 3; SI, Table S4). Due to the correlation between functional properties of agonists and receptor density (Kenakin et al., 2012), dimensionless efficacy to potency ratios relative to the reference agonist adrenaline (Δlog(E_max_/EC_50_)) were calculated (SI, Table S4). The calculated Δlog(E_max_/EC_50_) values suggested that changes of pEC_50_ and E_max_ were relevant for the partial agonists – salbutamol at β_1,2_AR and noradrenaline at β_2_AR.

## Discussion

To evaluate the mG_s_ sensor suitability for β_1,2_ARs, standard agonists and antagonists were functionally characterised using HEK293T cells stably co-expressing NlucN-mG_s_ and β_1_AR-NlucC or β_2_AR-NlucC fusion proteins. Overall, observed potencies were within the range of literature binding affinities (p*K*_i_) determined by radioligand displacement and pEC_50_ values obtained in functional cAMP accumulation assays (Baker, 2010; Hoffmann et al., 2004; Isogaya et al., 1999; Mistry et al., 2013). One obvious difference was carvedilol, where the pEC_50_ determined at β_1_AR differed from the reported p*K*_i_ by about two orders of magnitude (Hoffmann et al., 2004). In fact, this observation was consistent with previous findings of a second binding site at the β_1_AR, to which carvedilol and other antagonists bind with less affinity but increased efficacy (Baker et al., 2003; Baker et al., 2013; Konkar et al., 2000). On the other hand, E_max_ values of the partial agonists were lower in mG_s_ recruitment assays than in the more distal cAMP accumulation assays in the literature. Since the signal amplification along the signalling cascade potentially complicates the discrimination between full and partial agonists in drug discovery by leftward shift of pEC_50_ values and concomitant increase in E_max_, the presented mG_s_ sensors should be an attractive addition to future characterization of β_1,2_AR agonists, and thus might prevent overestimation of agonist intrinsic activities (Black and Leff, 1983; Kenakin, 1986; Thomsen et al., 2005).

The primary research questions were whether mini-G proteins can monitor GPCR activation at the low, ‘endogenous’ receptor level and whether β_1_AR is functionally expressed in addition to β_2_AR in HEK293T cells, which could lead to problems in ligand screening for receptor specificity. In our downstream assays using only native receptors in wild type HEK293T cells (cAMP accumulation and dynamic mass redistribution), the β-adrenergic ligand profiles did not match a single receptor response, confirming the presence of more than one receptor. Also, the noncompetitive effect observed for both antagonists was probably caused by low receptor reserve in the cell system with the native receptors, further confounding results (Kenakin, 2019).

In order to correctly ascertain the response of each βAR subtype in the system with these HEK293T cells, tagging was required. Our method of CRISPR/Cas9 tagging was cost-effective, utilising general cloning techniques with oligonucleotides and plasmid DNA. This method generated CRISPR/Cas9-modified HEK293T cells expressing β_1,2_AR-NlucC fusion proteins under endogenous promotion and could measure specific mG_s_ recruitment in response to agonists. Signal-to-baseline ratios were significantly lower at endogenously promoted β_1,2_AR-NlucC than with overexpressed receptors, as expected, but luminescent traces revealed similar kinetics. Overall, mG_s_ sensors revealed specific agonist modalities and thus should be well suited for the functional characterization of ligands at ‘endogenous’ receptor density. However, Δlog(E_max_/EC_50_) values significantly differed for partial agonists and further experiments would be needed for explanation. At low receptor density, the ratio of NlucC-labelled and natively expressed receptors might be relevant, especially for weaker assay responses elicited by partial agonists. Therefore, it would be prudent to quantify the absolute receptor expression, perhaps by radioligand saturation binding (unfortunately limited by the availability of selective radioligands for the β_1,2_AR).

To the best of our knowledge, we are the first to use the CRISPR/Cas9 system to replace a G_α_ protein subunit with a minimal G protein subunit (here, the G_αs_ protein with NlucN-mG_s_). When this system was tested with overexpressed β_1,2_AR-NlucCs, there appeared to be receptor- and ligand-specific changes in the pharmacological parameters. Receptor density is the probable cause for a reduced S/B ratio with the β_2_AR but not β_1_AR; our basal level of β_2_AR is higher than β_1_AR in our cells and overexpressed system, which could cause sequestering of the limited mG_s_ sensor. This theory is supported by the transient peak in the kinetic trace for the β_2_AR/CRISPR NlucN-mG_s_, and rightward potency shift and decreased E_max_ with the compounds. For β_1_AR, there was only a decrease in potency for adrenaline and salbutamol (no response). Both ligands have a higher potency for the native β_2_AR, which are already expressed natively in the cells and will sequester the mG_s_ sensor when activated. We therefore conclude that replacement of native effector proteins with synthetic constructs under the same promotor must be used with caution in order to retain receptor:sensor ratios useful to the biological question being posed.

Tagged, endogenously promoted β_2_AR have already been utilised for several purposes, such as deciphering the interaction between β_2_AR and the vascular endothelial growth factor receptor, assessing the binding properties of a fluorescent ligand, and monitoring receptor trafficking using fluorescence correlation spectroscopy (Goulding et al., 2021a; Goulding et al., 2021b; Kilpatrick et al., 2019). Here, for the first time, we have proved that the β_1_AR can be similarly tagged and used in HEK293T cells expressing both receptors to analyse intracellular responses to selective ligands. We anticipate that this knowledge will be useful to any future studies studying the β-adrenergic receptor subtypes in relation to one another.

Overall, with this study we confirm that native receptor populations are important to and directly affect recombinant cell line biology. These receptors and their effectors can be utilised to measure responses to the extracellular environment by specific tagging with CRISPR/Cas9 methods.

## Materials and Methods

### Materials

Dulbecco’s modified Eagle’s medium (DMEM) was purchased from Sigma-Aldrich (Taufkirchen, Germany) and Leibovitz’ L-15 medium (L-15) from Fisher Scientific (Nidderau, Germany). Foetal calf serum (FCS) and trypsin/EDTA were from Merck Biochrom (Darmstadt, Germany), coelenterazine h (5 mM in methanol, stored at −80 °C) was from BioSynth s.r.o (Bratislava, Slovakia) and furimazine was from Promega (Mannheim, Germany). The pcDNA3.1 vector was from Thermo Scientific (Nidderau, Germany) and the pIRES_puro_3 vector was a kindly provided by Prof. Dr. Gunter Meister (University of Regensburg, Regensburg, Germany). The pU6-SacB-scRNA-Cas9-T2A-mCherry plasmid was a gift from Kim Failor (Addgene plasmid # 117070) and pFETCh_Donor (EMM0021) was a gift from Eric Mendenhall & Richard M. Myers (Addgene plasmid # 63934). The cDNA encoding the β_1_AR was a kind gift from Ulrike Zabel (University of Würzburg, Würzburg, Germany) and the β_2_AR cDNA was purchased from the Missouri cDNA resource center [sic] (Rolla, MO, USA). Restriction enzymes *Bbs*I and *Bsa*I were from ThermoFisher (Braunschweig, Germany) and *Dpn*I, *Hin*dIII, *Xba*I, T4 DNA ligase, T7 endonuclease I and the NEBuilder® HiFi DNA Assembly Cloning Kit were from New England Biolabs (Frankfurt a. M., Germany). L-adrenaline (adren), L-noradrenaline tartrate (noradren), isoprenaline hydrochloride (isopren), salbutamol hemisulphate (salbut), carvedilol (carv), ICI 118551 hydrochloride (ICI118551) were from Sigma Aldrich. Xamoterol hemifumarate (xamo) was from Tocris Bioscience (Bristol, UK). Structures of the investigated ligands are given in the Supplementary Information (SI, Fig. S1). For preparation of stock solutions, the ligands were preferably dissolved in Millipore water. Adrenaline was dissolved in a mixture of 10 mM HCl and MDSO (50:50), and carvedilol was dissolved in DMSO.

### Isolation of Genomic DNA and Total RNA

Genomic DNA (gDNA) and total RNA were isolated from wildtype or CRISPR/Cas9 genome edited HEK293T cells. For gDNA isolation, the Monarch Genomic DNA Purification kit (New England Biolabs) was utilized with 1-2 x 10^6^ cells. RNA was isolated from 1-2 x 10^6^ cells using the Jena Bioscience Total RNA Purification kit (Jena Bioscience, Jena, Germany). Afterwards, co-precipitated DNA was removed by DNAse I (New England Biolabs) digest at 37°C for 10 min followed by a heat inactivation step at 75°C for 10 min. Then, RNA was purified from the DNAse reaction mix using the spin column RNA cleanup step of the Qiagen RNeasy Mini kit (Qiagen, Hilden, Germany) and reversely transcribed into cDNA using the ProtoScript II NEB cDNA synthesis kit (New England Biolabs).

### Generation of pcDNA3.1_β1,2AR-NlucC

The molecular cloning strategy of pIRES_puro_3 plasmids encoding the (LgBiT-) NlucN-mG_s_, fusion protein was described previously (Höring et al., 2020). For construction of pcDNA3.1 plasmids encoding the β_2_AR-NlucC or β_1_AR-NlucC fusion protein, a Gibson assembly protocol was performed. pcDNA3.1 encoding the small fragment of the NanoLuc (NlucC; VTGYRLFEEIL, ‘smBiT’) was linearized by restriction digest using *Hin*dIII and *Xba*I and then fused to the *Dpn*I digested PCR product of β_2_AR or β_1_AR, which contained respective overlaps to the vector backbone, using the NEBuilder® HiFi DNA Assembly Cloning Kit. All plasmid DNA was quantified by UV-Vis absorbance using a NanoDrop spectrophotometer (ThermoFisher). All sequences were verified by sequencing performed by Eurofins Genomics (Eurofins Genomics LLC, Ebersberg, Germany).

### Generation of pU6_gRNA-Cas9-T2A-mCherry Plasmids

For gRNA and Cas9 expression, the pU6_SacB-Cas9-T2A-mCherry plasmid was used. The GNAS gene coding the G_αs_ subunit was targeted using a gRNA (CTACAACATGGTCATCCGGG) previously described by Stallaert et al. (2017) to generate G_αs_ knockout cells. An overview of primer sequences used for the generation and verification of HEK293T_CRISPR_β_1,2_AR-NlucC cells is given in the Supplementary Information (SI, Table S1). Using the Benchling web server (www.benchling.com), oligonucleotides encoding the gRNA (oligos; β_1_AR: O1/O2, β_2_AR: O3/O4; SI, Table S1), were selected upstream from the *Streptococcus pyogenes* Cas9 (spCas9) PAM (β_1_AR-PAM: GGG, β_2_AR-PAM: AGG) and within 50 bp of the STOP codon of the receptor. Briefly, the single stranded gRNA oligos were annealed in NEBuffer 2 (New England Biolabs) at 95 °C for 5 min and left to cool. Thereafter, each of the annealed gRNA oligos were subcloned into the pU6_SacB-scRNA-Cas9-mCherry vector using a Golden Gate assembly protocol with *Bbs*I and T4 DNA ligase. Plasmids were transformed into TOP10’F *E. coli* (made competent in-house) and extracted in maxipreps (Qiagen).

Cas9 cleavage of β_1,2_AR target sites were verified by T7 endonuclease digests recognising DNA mismatches. For this purpose, 600,000 HEK293 cells were transiently transfected with 2 µg of plasmid DNA using XtremeGene HP transfection reagent (ratio 1 μg DNA : 3 μL XtremeGene HP) and the gDNA was extracted after 48 hours. Diagnostic DNA templates were amplified from gDNA using primer pairs O5/O6 for the β_1_AR and O7/O8 for the β_2_AR (SI Table S1) and digested using T7 endonuclease I according to the supplier’s protocol. Digested PCR products were visually inspected after separation by 0.8% agarose gel electrophoresis and revealed cleaved PCR products, which indicated DNA mismatches caused by Cas9 nuclease cleavage (data not shown).

### Generation of Homology Directed Repair Template Plasmids

To fuse NlucC to the C-terminus of endogenously expressed β_2_AR and β_1_AR, the CRISPR /Cas9 protocol was adapted from the Mendenhall and Myers labs’ FLAG tagging approach (Savic et al., 2015). For homology directed repair (HDR), a donor plasmid (pFETCh_Donor) was used. In a first cloning step, the FLAG sequence of the pFETCh vector was replaced by the NlucC sequence. Primers O9/O10 (SI, Table S1) were used to linearise the vector and corresponding overlaps were attached to the NlucC sequence using primers O11/O12. Standard Gibson assembly cloning was then used to generate the pFETCh_NlucC. Sequences of the homology arms 1 and 2 (HOM1, HOM2) specific for β_1_AR and β_2_AR were determined using the Benchling (www.benchling.com) and Ensembl (www.ensembl.org) web servers. HOM1 comprised the C-terminal end of either the β_1_AR or β_2_AR (without the stop codon) and HOM2 was selected downstream of the receptor sequence in the genomic DNA. Specifically, homology arm primers were selected to remove the native β_1,2_AR stop codons as well as the spCas9 PAM to prevent further enzymatic cleavage after successful HDR (β_1_AR-PAM: GGG to GTG, β_2_AR-PAM: AGG to AGA). HOM1 and HOM2 were amplified from HEK293T gDNA and overlaps for type IIS enzyme digest (*Bsa*I for HOM1 and *Bbs*I for HOM2) were added by PCR. For the β_1_AR, the primer pairs O13/O14 and O15/O16, and for the β_2_AR the primer pairs O17/O18 and O19/O20 were used to obtain HOM1 and HOM2, respectively (SI, Table S1). The pFETCh_NlucC plasmid was digested with *Bbs*I and *Bsa*I and annealed with purified HOM1 and HOM2 PCR products in a Gibson assembly reaction yielding pFETCh_HOM1-NlucC-P2A-NeoR-HOM2 (pFETCh_β_1,2_AR-NlucC) specific for either β_1_AR or β_2_AR.

For expression of NlucN-mG_s_ under the GNAS promotor, pFETCh_NlucN-mG_s_ was first generated using the linearised vector (pFETCh_Donor PCR with O9 and O10) and NlucN-mG_s_ with complementary ends (PCR using O23/O24, SI, Table S1) in a standard Gibson Assembly reaction. Homology arms for the area of the GNAS gene around the Cas9 cut site were taken from HEK293T gDNA in a PCR reaction using primer pairs O25/O26 and O27/O28, respectively. The same reaction as for β_1,2_AR homology arm addition was then used to generate pFETCh_HOM1-NlucN-mG_s_-P2A-NeoR-HOM2 (pFETCh_NlucN-mG_s_).

Homology arm and editing sequences for all constructs are provided in the Supplementary Information (SI, Table S3).

### Cell Culture

HEK293T cells (a kind gift from Prof. Dr. Wulf Schneider; Institute for Medical Microbiology and Hygiene, Regensburg, Germany), were cultured in DMEM supplemented with 10% FCS at 37 °C in humidified, 5% CO_2_ atmosphere. The cells were periodically tested negative for mycoplasma contamination using the Venor GeM Mycoplasma Detection Kit (Minerva Biolabs, Berlin, Germany).

### Generation of Stable Transfectants

To generate HEK293T cells stably co-expressing NlucN-mG_s_ and β_2_AR-NlucC or β_1_AR-NlucC (overexpressed), the parental HEK293T_NlucN-mG_s_ cells (Höring et al., 2020) were seeded into a 6-well cell culture plate (Sarstedt, Nümbrecht, Germany) at a cell density of 0.3 x 10^6^ cells.mL^-1^ (2 mL per well) and allowed to attach overnight. The next day, 2 μg of pcDNA3.1_β_1,2_AR-NlucC plasmid DNA was transfected into the cells using the XtremeGene HP transfection protocol (ratio 1 μg DNA: 3 μL XtremeGene HP). Finally, the cells were then cultured in DMEM supplemented with 10% FCS, 1 μg.mL^-1^ puromycin and 600 μg.mL^-1^ G418 for sustained selection pressure.

To generate genome edited HEK293T cells (HEK293T_CRISPR_β_1,2_AR-NlucC or HEK293T_CRISPR_NlucN-mG_s_), HEK293T wildtype cells were seeded in a T25 cell culture flask at a cell density of 0.3 x 10^6^ cells.mL^-1^. After the cells had grown overnight, 2 μg of the pFETCh_β_1_AR, pFETCh_β_2_AR or pFETCh_NlucN-mGs and 1 μg of the corresponding pU6_gRNA-Cas9-T2A-mCherry plasmids were transfected using 12 μL XtremeGene HP transfection reagent (1:4 ratio). After two weeks of antibiotic selection using 1000 μg.mL^-1^ G418, single clones were isolated from the heterogenous cell population by dilution cloning. Afterwards, single clones were cultured in DMEM (full medium) containing 600 μg.mL^-1^ G418.

### Transient Transfections

To verify successful generation of β_1,2_AR-NlucC using CRISPR/Cas9, pooled HEK293T CRISPR β_1,2_AR-NlucC cells were seeded onto a 6-well cell culture plate at a density of 0.3 x 10^6^ cells.mL^-1^ (2 mL per well) and were transiently transfected the next day with a total amount of 1 μg plasmid DNA using 3 μL XtremeGene HP (1:3 ratio). For this purpose, different mixtures were prepared comprising 17 ng, 35 ng, 70 ng, 140 ng, 280 ng or 567 ng of pIRES_puro_3 NlucN-mG_s_ plasmid DNA in combination with 983 ng, 965 ng, 930 ng, 860 ng, 720 ng or 433 ng of an empty pIRES_puro_3 plasmid (mock DNA), respectively. Transfectants were incubated at 37 °C for 48 h in a humidified, 5% CO_2_ atmosphere.

To screen single clones isolated from pooled HEK293T_CRISPR_β_1,2_AR-NlucC or HEK293T_CRISPR_NlucN-mG_s_ cells, the single clones were seeded in duplicate to 96-well cell culture plates at a density of 0.3 x 10^6^ cells.mL^-1^ (100 μL per well) and transiently transfected with pIRES_puro_3_NlucN-mG_s_ or p3.1_ β_2_AR-NlucC plasmid DNA using XtremeGene HP transfection reagent (1:3 ratio). For this, a transfection mixture was prepared in 250 μL DMEM containing 2600 ng plasmid DNA and 7.8 μL XtremeGene HP. After 15 min incubation, the transfection mix was diluted to 1000 μL and 100 μL each of the diluted transfection complex were added to the cells in the 96-well plate. Transfectants were incubated at 37 °C for 48 h in a humidified, 5% CO_2_ atmosphere.

For HEK293T_CRISPR_NlucN-mG_s_, an extra step to decide the amount of receptor plasmid DNA using a transient transfection with differing amounts of receptor-NlucC plasmid DNA (250 ng, 500 ng and 1000 ng) made up to 1000 ng with pcDNA3.1 empty plasmid, using the same protocol as follows.

To functionally characterize β_1,2_AR ligands, CRISPR/Cas9 positive single clones, HEK293T_CRISPR_β_1_AR-NlucC clone 11, HEK293T_CRISPR_β_2_AR-NlucC clone 35 or HEK293T_CRISPR_NlucN-mG_s_ clone 83 were seeded into 6-well cell culture plates at a cell density of 0.3 x 10^6^ cells.mL^-1^ (2 mL per well) and allowed to attach overnight. The next day, the cells were transfected with a total plasmid DNA amount of 1 μg using 3 μL XtremeGeneHP transfection reagent (1:3 ratio). For CRISPR_β_1,2_AR cells, 280 ng of pIRES_puro_3_NlucN-mG_s_ and 720 ng of empty pIRES_puro_3 (mock) plasmid DNA were used. For CRISPR_ mG_s_ cells, 500 ng of p3.1_β_1,2_AR-NlucC and 500 ng of empty pcDNA3.1 (mock) plasmid DNA were used. Transfectants were incubated at 37 °C for 48 h in a humidified, 5% CO_2_ atmosphere.

### Mini-G Protein Recruitment Assays

The day prior to the experiment, stable or transient transfectants (except for single clone screens) were detached using trypsin (0.05% trypsin, 0.02% EDTA in PBS) and centrifuged (700 g, 5 min). Thereafter, the cells were resuspended in 2.5 mL L-15 supplemented with 10 mM HEPES (Serva, Heidelberg, Germany) and 5% FCS (assay medium) and 80 μL or 40 μL per well were seeded into white flat-bottom 96- or 384-well microtiter plates, respectively (Brand, Wertheim, Germany). For single clone screens, the cell culture medium (DMEM, full medium) was removed from the cells, which had been transiently transfected in the 96-well plate. Following a washing step, 80 μL of assay medium were added to the cells. All cells were then incubated overnight at 37 °C in a water-saturated atmosphere without additional CO_2_.

For mini-G protein recruitment assays performed in 96-well plates, the assay protocol was as previously described (Höring et al., 2020). When assays were performed in 384-well plates, the protocol was adjusted as follows: The furimazine substrate was diluted 1:500 in L-15 and 5 μL was added to the cells (final assay dilution: 1:5000). The plate was transferred to the pre-heated (37 °C) Tecan Infinite 200 Pro plate reader (Tecan, Männedorf, Switzerland) and 5 plate repeats of basal luminescence were recorded (∼16.5 min). Thereafter, 5 μL of the agonist serial dilutions prepared in L-15 were added to each well (final assay volume: 50 μL) and luminescent signals were recorded for an additional 15 plate repeats (∼49.5 min). An integration time of 0.2 s per well was used to capture the luminescence and plate repeats lasted 198 s.

### CAMYEN cAMP Assay

The CAMYEN BRET-based assay was used to determine cAMP generation by β_1,2_AR ligands with native receptors. HEK293T-CAMYEN stable cell lines (cultured with 600 ng.mL^-1^ G418) were used as described in a previously reported protocol (Gleixner et al., 2023), with minor alterations described as follows. Cells without transfected receptor plasmid DNA were plated in opaque, white 96-well plates instead of 384-well plates. Each well contained 75,000 cells in 80 µL L-15 media with 10 mM HEPES and 5% FCS. On the day of the experiment, cells were incubated in 10 µL.well^-1^ of 5 µM coelenterazine h without forskolin but including 100 µM 3-isobutyl-1-methylxanthine (IBMX, diluted in DMSO). The baseline luminescence was then recorded for both “Blue”(< 470nm) and “Green” (520−580 nm) filter wavelengths on a preheated (37 °C) Tecan Infinite Lumi at an integration time of 100 ms for 30 min (12 cycles). Agonists, prepared at 10x concentration, were added after the baseline read at 10 µL.well^-1^. Control wells added 10 µM forskolin or buffer. For antagonists, the antagonist was added at the same time as the substrate (30 min preincubation) and diluted isoprenaline was then added after the baseline read. The plate was then read for a further 60 min (23 cycles).

### Dynamic Mass Redistribution Assay

Label-free analysis of ligand innovation in HEK293T wild type cells was performed with the dynamic mass redistribution assay as previously described (Seibel-Ehlert et al., 2021) – adapted from Schröder et al. (2011) – with minor modifications. No antibiotics were used with the HEK293T wildtype cells at any stage of culturing or the assay. As previously performed, cells were used at 1 × 10^6^ cells.mL^-1^ with 90 µL.well^-1^ in an assay-specific, label-free 96 well plate (Cat. No. 5080, Corning Life Sciences, Amsterdam, Netherlands). Unlike previously described, the plates were precoated with poly-D-lysine (mol. wt 70,000-150,000, Sigma-Aldrich, Taufkirchen, Germany). The same assay media, Hank’s Balanced Salt Solution (HBSS, Gibco, Thermo Fisher) with 20 mM HEPES, was used in these experiments.

### Data Analysis

Data were analysed using GraphPad Prism 9 software (San Diego, CA, USA). In a first baseline-correction step, the relative luminescence units (RLU) or BRET ratios (520−580 nm / < 470 nm wavelength recordings) were corrected for slight inter-well variations caused by differences in cell density and substrate concentration by dividing the data of each well by its luminescence prior to the addition of the ligands. In a second baseline-correction step, all data were divided by the mean luminescence or BRET ratio of L-15 responses, producing relative increases in luminescence. For each well, the area under curve (AUC) of the traces was normalized to the maximum response of the reference agonist (10 μM adren) and L-15 (0% control). For assays performed in 96-well plates using an EnSpire plate reader (Perkin Elmer, Rodgau, Germany), AUCs were calculated from signals of 45 min (90 plate repeats of 30 s length each), or for the Tecan Infinite Lumi, 60 min (23 cycles of approximately 156 s each). For assays performed in 384-well plates, AUCs were calculated of signals within 49.5±0.05 min (15 plate repeats of 198 s length each). Normalized data were then fitted against the logarithmic ligand concentrations with variable slope (log(c) vs. response – variable slope (four parameters)) to generate pEC_50_ and E_max_ values, unless otherwise stated.

### Calculation of Δlog(Emax/EC50)

For each experiment, the efficacy to potency ratio E_max_/EC_50_ was determined and logarithmically transformed into log(E_max_/EC_50_). Subsequently, Δlog(E_max_/EC_50_) for each agonist (A) were calculated using the means ± SEM of single log(E_max_/EC_50_) values relative to the reference agonist adrenaline (ref) as follows (Griffin et al., 2007; Kenakin et al., 2012):

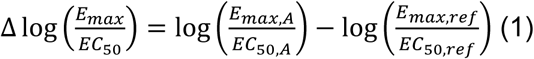

For Δlog(E_max_/EC_50_) values, error propagation was performed according to the following equation:

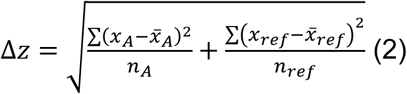

where 𝑥 is the function of Δlog(E_max_/EC_50_) of each agonist (A) and the reference agonist adrenaline (ref). Consequently, Δ𝑧 represents the propagated error of z.

### Sample Sizes and Statistics

Throughout the manuscript, the sample size *n* was defined as a single experiment performed on a separate day with a different cell passage and separate transient transfection, where applicable. Statistics were only performed on data where *n* ≥ 5.

## Acknowledgements

We would like to acknowledge our undergraduate placement students Annalena Schiestl and Sebastian Klinger and our colleagues Denise Mönnich and Maria Beer-Kroen for their help in optimising the assays used in this manuscript. We also thank Professor Pierre Koch and Dr Max Keller for providing the infrastructure to carry out experiments and the Deutsche Forschungsgemeinschaft for funding this project.

## Data Availability

All relevant data can be found within the article and its supplementary information. Raw data is available upon reasonable request.

## Abbreviations

adren: L-adrenaline
AUC: area under the curve
βAR: β-adrenoceptor
BRET: Bioluminescence resonance energy transfer
Cas: CRISPR-associated protein
carv: carvedilol
CRISPR: clustered regularly interspaced short palindromic repeats
FCS: foetal calf serum
GPCR: G protein coupled receptor
HDR: homology directed repair
ICI: or ICI118551 ICI 118551 hydrochloride
isopren: isoprenaline hydrochloride
gDNA: genomic DNA
gRNA: guide RNA
mG_s_: minimal or ‘mini’ G_αs_ protein
NanoLuc/Nluc: Nanoluciferase
noradren: L-noradrenaline tartrate
PAM: protospacer adjacent motif
RLU: relative luminescence units
salbut: salbutamol hemisulphate
SI: supplementary information
SLC: split luciferase complementation
spCas9: *Streptococcus pyogenes* Cas9
xamo: xamoterol hemifumarate

