## Supplemental Information for "Endogenous Promotor-Driven Split Nanoluciferase Biosensor for Assessing G Protein Recruitment"

^#^previous affiliation

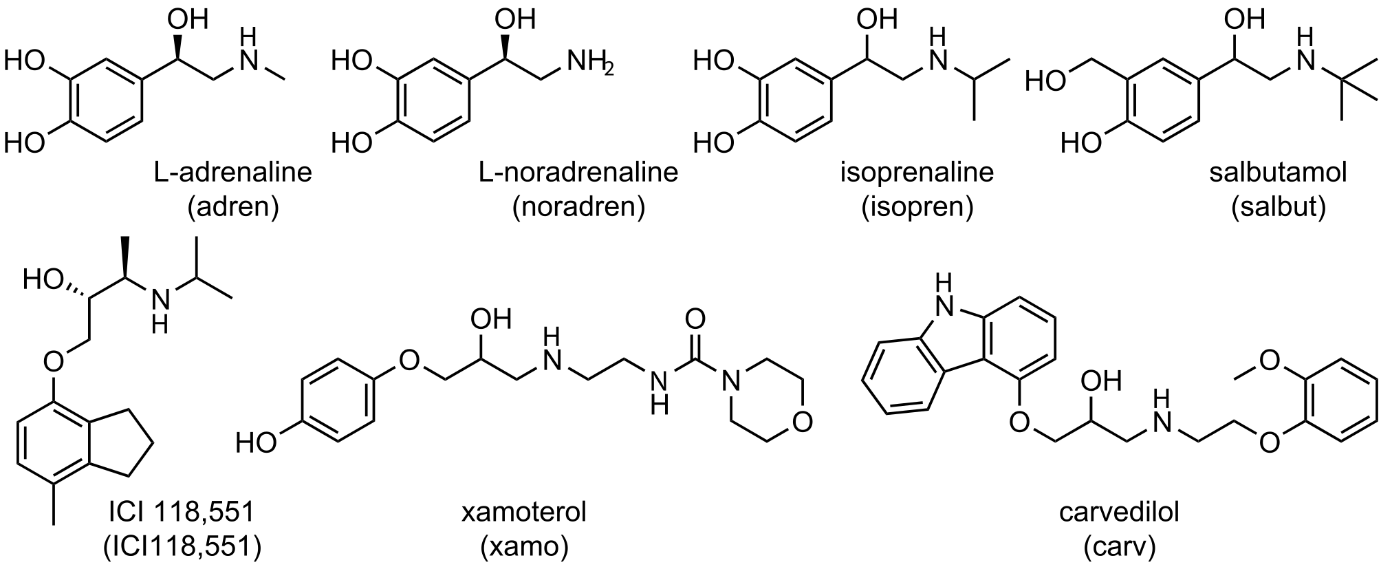

**Figure S1** – Structures of investigated β_1,2_AR ligands.

**Table S1** – Oligonucleotide sequences used in the manuscript.

| Oligo name | | Sequence |
| --- | --- | --- |
| O1 | β_1_AR gRNA fw | CACCGCCTCGGAATCCAAGGTGTA |
| O2 | β_1_AR gRNA rv | AAACTACACCTTGGATTCCGAGGC |
| O3 | β_2_AR gRNA fw | CACCGATAACATTGATTCACAAGGG |
| O4 | β_2_AR gRNA rv | AAACCCCTTGTGAATCAATGTTATC |
| O5 | β_1_AR diag fw | CTGCTACAACGACCCCAAGT |
| O6 | β_1_AR diag rv | TCCCCTAACCCACCCATCTT |
| O7 | β_2_AR diag fw | TCCCCTTATCTACTGCCGGA |
| O8 | β_2_AR diag rv | AACAGGTGCAATGAAGGCAT |
| O9 | pFETCh(-)FLAG fw | CGCTCAGAGACCCGCGTAAG |
| O10 | pFETCh(-)FLAG rv | GTTTCAGGAAGCGGAGCTAC |
| O11 | NlucC fw | AGTTAGTAGCTCCGCTTCCTGAAACGAGAATCTCCTCGAACAGCCG |
| O12 | NlucC rv | GAGACCTTACGCGGGTCTCTGAGCGGAGGAGGTGGCGGATCCGGT |
| O13 | β_1_AR HOM1 fw | TCCCCGACCTGCAGCCCAGCTTGCTACAACGACCCCAAG |
| O14 | β_1_AR HOM1 rv | ATCCGCCACCTCCTCCGCTCCCCACCTTGGATTCCGAGGC |
| O15 | β_1_AR HOM2 fw | AGTTCTTCTGATTCGAACATCTAGCTGCCCGGCGCGGGGC |
| O16 | β_1_AR HOM2 rv | TGGAGAGGACTTTCCAAGCCCAGGCGCGCGGGGGAC |
| O17 | β_2_AR HOM1 fw | TCCCCGACCTGCAGCCCAGCTAGAATAAGGCCCGGGTGATCATTCT |
| O18 | β_2_AR HOM1 rv | ATCCGCCACCTCCTCCGCTCCCCAGCAGTGAGTCATTTGTACTACAATTT |
| O19 | β_2_AR HOM2 fw | AGTTCTTCTGATTCGAACATCTAAAGCAGTTTTTCTACT |
| O20 | β_2_AR HOM2 rv | TGGAGAGGACTTTCCAAGAGACTCAAAGGCAAATGA |
| O21 | β_2_AR CRISPR control fw | AAGCTGCTCCTCAAATCCCT |
| O22 | β_2_AR CRISPR control rv | CCTTACCTCCTTCTTGCCCA |
| O23 | NlucN-mG_s_ fw | AGTTAGTAGCTCCGCTTCCTGAAACCAACAGCTCGTATTGCCG |
| O24 | NlucN-mG_s_ rv | GAGACCTTACGCGGGTCTCTGAGCGATGGTGTTCACCCTGGAG |
| O25 | GNAS HOM1 fw | TCCCCGACCTGCAGCCCAGCTACCATCGCTGTGTAGTGTGG |
| O26 | GNAS HOM1 rv | CGAAGTCCTCCAGGGTGAACACCATGTTCCCGAGGCAGCCCAAG |
| O27 | GNAS HOM2 fw | AGTTCTTCTGATTCGAACATCAGTAAGACCGAGGACCAG |
| O28 | GNAS HOM2 rv | TGGAGAGGACTTTCCAAGCCTTTACCGCTCCGATCCTT |

**Table S2** – Potencies (pEC_50_, p*K*_b_) and efficacies (E_max_) of selected standard agonists and antagonists obtained in mini-G protein recruitment assays using HEK293T cells co-expressing NlucN-mG_s_ and β_1,2_AR-NlucC fusion proteins. Responses were normalized to L-15 as solvent control (0%) and to the maximal response elicited by 100 µM (β_1_AR) or 10 µM adren (β_2_AR) for agonists and 100 nM adren for antagonists (100%). Data represent means ± SEM from at least three independent experiments (*n* ≥ 3), performed in triplicate.

|  |  | Mini-G Protein Recruitment | | cAMP Accumulation | | Competition  Binding |
| --- | --- | --- | --- | --- | --- | --- |
| Subtype | Cpd. | pEC_50_ (p*K*_b_)± SEM | E_max_ (%) ± SEM | pEC_50_ | E_max_ (%) | p*K*_i_ |
| β_1_AR | adren | 6.86 ± 0.02 | 100 | 7.61^a^ | 102^a^ | 5.40^c^ |
|  | noradren | 6.21 ± 0.06 | 115 ± 16 | 7.94^a^ | 102^a^ | 5.45^c^ |
|  | isopren | 7.32 ± 0.03 | 110± 1* | 8.59^a^ | 100^a^ | 6.65^c^ |
|  | salbut | 5.40 ± 0.02 | 37 ± 1 | 6.21^a^ | 104^a^ | 5.61^c^ |
|  | xamo | 7.21 ± 0.14 | 13 ± 1 | 7.96^a^ | 48^a^ | 7.00^e^ |
|  | ICI118,551 | (6.38) ± 0.03 | n.d. |  |  | 7.31^c^ |
|  | carv | 6.72 ± 0.16 | 6.38 ± 1.14 | 7.64^a^ | 10^a^ | 9.23^c^ |
| β_2_AR | adren | 7.55 ± 0.06 | 100 | 7.93^b^ | 102^b^ | 6.13^d^ |
|  | noradren | 5.12 ± 0.05 | 85 ± 2 | 6.36^b^ | 103^b^ | 4.58^d^ |
|  | isopren | 7.21 ± 0.12 | 107 ± 5 | 8.22^b^ | 100^b^ | 6.34^d^ |
|  | salbut | 7.15 ± 0.09 | 82 ± 1 | 7.72^b^ | 96^b^ | 5.66^d^ |
|  | xamo | (6.09) ± 0.06 | n.d. | 6.15^b^ | 5^b^ | 5.85^f^ |
|  | ICI118,551 | (9.09) ± 0.14 | n.d. |  |  | 9.16^d^ |
|  | carv | (8.29) ± 0.03 | n.d. |  |  | 8.96^d^ |

Reference data were taken from: Functional [^3^H]cAMP accumulation assays using CHO K1 cells expressing ^a^β_1_AR or ^b^β_2_AR (data normalized to maximal responses of isoprenaline) (Baker et al., 2013; Mistry et al., 2013) and (-)-3-[^125^I]-iodocyanopindolol displacement assays using CHO cells expressing ^c^β_1_AR or ^d^β_2_AR (Hoffmann et al., 2004) or COS-7 cells expressing ^e^β_1_AR or ^f^β_2_AR (Isogaya et al., 1999).

**Table S3** – Homology and editing sequences for CRISPR/Cas9 modified cells with human genome. CAPS = homology arms, **bold + underlined** = fused protein, underlined = P2A site and neomycin resistance gene.

| **Gene and edit** | **Sequence** |
| --- | --- |
| *ADRB1*-NlucC | TGCTACAACGACCCCAAGTGCTGCGACTTCGTCACCAACCGGGCCTACGCCATCGCCTCGTCCGTAGTCTCCTTCTACGTGCCCCTGTGCATCATGGCCTTCGTGTACCTGCGGGTGTTCCGCGAGGCCCAGAAGCAGGTGAAGAAGATCGACAGCTGCGAGCGCCGTTTCCTCGGCGGCCCAGCGCGGCCGCCCTCGCCCTCGCCCTCGCCCGTCCCCGCGCCCGCGCCGCCGCCCGGACCCCCGCGCCCCGCCGCCGCCGCCGCCACCGCCCCGCTGGCCAACGGGCGTGCGGGTAAGCGGCGGCCCTCGCGCCTCGTGGCCCTGCGCGAGCAGAAGGCGCTCAAGACGCTGGGCATCATCATGGGCGTCTTCACGCTCTGCTGGCTGCCCTTCTTCCTGGCCAACGTGGTGAAGGCCTTCCACCGCGAGCTGGTGCCCGACCGCCTCTTCGTCTTCTTCAACTGGCTGGGCTACGCCAACTCGGCCTTCAACCCCATCATCTACTGCCGCAGCCCCGACTTCCGCAAGGCCTTCCAGGGACTGCTCTGCTGCGCGCGCAGGGCTGCCCGCCGGCGCCACGCGACCCACGGAGACCGGCCGCGCGCCTCGGGCTGTCTGGCCCGGCCCGGACCCCCGCCATCGCCCGGGGCCGCCTCGGACGACGACGACGACGATGTCGTCGGGGCCACGCCGCCCGCGCGCCTGCTGGAGCCCTGGGCCGGCTGCAACGGCGGGGCGGCGGCGGACAGCGACTCGAGCCTGGACGAGCCGTGCCGCCCCGGCTTCGCCTCGGAATCCAAGGTG**gggagcggaggaggtggcggatccggtggtgggggcagtgtgaccggctaccggctgttcgaggagattctcgtttcaggaagcgga**gctactaacttcagcctgctgaagcaggctggagacgtggaggagaaccctggacctggatcgtttcgcatgattgaacaagatggattgcacgcaggttctccggccgcttgggtggagaggctattcggctatgactgggcacaacagacaatcggctgctctgatgccgccgtgttccggctgtcagcgcaggggcgcccggttctttttgtcaagaccgacctgtccggtgccctgaatgaactgcaggacgaggcagcgcggctatcgtggctggccacgacgggcgttccttgcgcagctgtgctcgacgttgtcactgaagcgggaagggactggctgctattgggcgaagtgccggggcaggatctcctgtcatctcaccttgctcctgccgagaaagtatccatcatggctgatgcaatgcggcggctgcatacgcttgatccggctacctgcccattcgaccaccaagcgaaacatcgcatcgagcgagcacgtactcggatggaagccggtcttgtcgatcaggatgatctggacgaagagcatcaggggctcgcgccagccgaactgttcgccaggctcaaggcgcgcatgcccgacggcgaggatctcgtcgtgacccatggcgatgcctgcttgccgaatatcatggtggaaaatggccgcttttctggattcatcgactgtggccggctgggtgtggccgaccgctatcaggacatagcgttggctacccgtgatattgctgaagagcttggcggcgaatgggctgaccgcttcctcgtgctttacggtatcgccgctcccgattcgcagcgcatcgccttctatcgccttcttgacgagttcttctgattcgaacatcTAGTGCCCGGCGCGGGGCGCGGACTCCGGGCACGGCTTCCCAGGGGAACGAGGAGATCTGTGTTTACTTAAGACCGATAGCAGGTGAACTCGAAGCCCACAATCCTCGTCTGAATCATCCGAGGCAAAGAGAAAAGCCACGGACCGTTGCACAAAAAGGAAAGTTTGGGAAGGGATGGGAGAGTGGCTTGCTGATGTTCCTTGTTGTTTTTTTTTTCTTTTCTTTTCTTTCTTCTTCTTTTTTTTTTTTTTTTTTTTTTCTGTTTGTGGTCCGGCCTTCTTTTGTGTGTGCGTGTGATGCATCTTTAGATTTTTTTCCCCCACCAGGTGGTTTTTGACACTCTCTGAGAGGACCGGAGTGGAAGATGGGTGGGTTAGGGGAAGGGAGAAGCATTAGGAGGGGATTAAAATCGATCATCGTGGCTCCCATCCCTTTCCCGGGAACAGGAACACACTACCAGCCAGAGAGAGGAGAATGACAGTTTGTCAAGACATATTTCCTTTTGCTTTCCAGAGAAATTTCATTTTAATTTCTAAGTAATGATTTCTGCTGTTATGAAAGCAAAGAGAAAGGATGGAGGCAAAATAAAAAAAAATCACGTTTCAAGAAATGTTAAGCTCTTCTTGGAACAAGCCCCACCTTGCTTTCCTTGTGTAGGGCAAACCCGCTGTCCCCCGCGCGCCTGGG |
| *ADRB2*-NlucC | AGAATAAGGCCCGGGTGATCATTCTGATGGTGTGGATTGTGTCAGGCCTTACCTCCTTCTTGCCCATTCAGATGCACTGGTACCGGGCCACCCACCAGGAAGCCATCAACTGCTATGCCAATGAGACCTGCTGTGACTTCTTCACGAACCAAGCCTATGCCATTGCCTCTTCCATCGTGTCCTTCTACGTTCCCCTGGTGATCATGGTCTTCGTCTACTCCAGGGTCTTTCAGGAGGCCAAAAGGCAGCTCCAGAAGATTGACAAATCTGAGGGCCGCTTCCATGTCCAGAACCTTAGCCAGGTGGAGCAGGATGGGCGGACGGGGCATGGACTCCGCAGATCTTCCAAGTTCTGCTTGAAGGAGCACAAAGCCCTCAAGACGTTAGGCATCATCATGGGCACTTTCACCCTCTGCTGGCTGCCCTTCTTCATCGTTAACATTGTGCATGTGATCCAGGATAACCTCATCCGTAAGGAAGTTTACATCCTCCTAAATTGGATAGGCTATGTCAATTCTGGTTTCAATCCCCTTATCTACTGCCGGAGCCCAGATTTCAGGATTGCCTTCCAGGAGCTTCTGTGCCTGCGCAGGTCTTCTTTGAAGGCCTATGGGAATGGCTACTCCAGCAACGGCAACACAGGGGAGCAGAGTGGATATCACGTGGAACAGGAGAAAGAAAATAAACTGCTGTGTGAAGACCTCCCAGGCACGGAAGACTTTGTGGGCCATCAAGGTACTGTGCCTAGCGATAACATTGATTCACAAGGGAGAAATTGTAGTACAAATGACTCACTGCTG**gggagcggaggaggtggcggtaccggtggtgggggcagtgtgaccggctaccggctgttcgaggagattctcgtttcaggaagcgga**gctactaacttcagcctgctgaagcaggctggagacgtggaggagaaccctggacctggatcgtttcgcatgattgaacaagatggattgcacgcaggttctccggccgcttgggtggagaggctattcggctatgactgggcacaacagacaatcggctgctctgatgccgccgtgttccggctgtcagcgcaggggcgcccggttctttttgtcaagaccgacctgtccggtgccctgaatgaactgcaggacgaggcagcgcggctatcgtggctggccacgacgggcgttccttgcgcagctgtgctcgacgttgtcactgaagcgggaagggactggctgctattgggcgaagtgccggggcaggatctcctgtcatctcaccttgctcctgccgagaaagtatccatcatggctgatgcaatgcggcggctgcatacgcttgatccggctacctgcccattcgaccaccaagcgaaacatcgcatcgagcgagcacgtactcggatggaagccggtcttgtcgatcaggatgatctggacgaagagcatcaggggctcgcgccagccgaactgttcgccaggctcaaggcgcgcatgcccgacggcgaggatctcgtcgtgacccatggcgatgcctgcttgccgaatatcatggtggaaaatggccgcttttctggattcatcgactgtggccggctgggtgtggccgaccgctatcaggacatagcgttggctacccgtgatattgctgaagagcttggcggcgaatgggctgaccgcttcctcgtgctttacggtatcgccgctcccgattcgcagcgcatcgccttctatcgccttcttgacgagttcttctgattcgaacatcTAAAGCAGTTTTTCTACTTTTAAAGACCCCCCCCCCCAACAGAACACTAAACAGACTATTTAACTTGAGGGTAATAAACTTAGAATAAAATTGTAAAATTGTATAGAGATATGCAGAAGGAAGGGCATCCTTCTGCCTTTTTTATTTTTTTAAGCTGTAAAAAGAGAGAAAACTTATTTGAGTGATTATTTGTTATTTGTACAGTTCAGTTCCTCTTTGCATGGAATTTGTAAGTTTATGTCTAAAGAGCTTTAGTCCTAGAGGACCTGAGTCTGCTATATTTTCATGACTTTTCCATGTATCTACCTCACTATTCAAGTATTAGGGGTAATATATTGCTGCTGGTAATTTGTATCTGAAGGAGATTTTCCTTCCTACACCCTTGGACTTGAGGATTTTGAGTATCTCGGACCTTTCAGCTGTGAACATGGACTCTTCCCCCACTCCTCTTATTTGCTCACACGGGGTATTTTAGGCAGGGATTTGAGGAGCAGCTTCAGTTGTTTTCCCGAGCAAAGTCTAAAGTTTACAGTAAATAAATTGTTTGACCATGCCTTCATTGCACCTGTTTCTCCAAAACCCCTTGACTGGAGTGCTGTTGCCTCCCCCACTGGAAACCGCAGGTAACTACTTGTAATTACTGCCCATGACTTAATGTAGAATGATACAAGAATGACATGCACAGATTGCTTAACCCTTTCATTTGCCTTTGAGTCT |
| *GNAS* KO, NlucN-mG_s_ | TGTGCCATTGACTTAGTGCTGCATAACTGTGGGACGGTCACTTCCGTTGAGCCTGACCTTGTAGAGAGACACAAATAGTTGGCAAATTGATGTGAGCGCTGTGAACACCCCACGTGTCTTTCTTTTTCTCCCAGCTTCCTGGACAAGATCGACGTGATCAAGCAGGCTGACTATGTGCCGAGCGATCAGGTGTGCAAAACCCCTCCCCACCAGAGGACTCTGAGCCCTCTTTCCAAACTACTCCAGACCTTTGCTTTAGATTGGCAATTATTACTGTTTCGGTTGGCTTTGGTGAGATCCATTGACCTCAATTTTGTTTCAGGACCTGCTTCGCTGCCGTGTCCTGACTTCTGGAATCTTTGAGACCAAGTTCCAGGTGGACAAAGTCAACTTCCAGTAAGCCAACTGTTACCTTTTTATATAACAGAGATCATGGTTTCTTGACATTCACCCCAGTCCCTCTGGAATAACCAGCTGTCCTCCTCCCCACCAGCATGTTTGACGTGGGTGGCCAGCGCGATGAACGCCGCAAGTGGATCCAGTGCTTCAACGGTAGGATGCTGTGGGCTTGGCTGTTCGTAAAGAACGCTTTGCTTCTGTGTTGTTAGGGATCAGGGTCGCTGCTCACGCTCTTGGCTTTGCTCTCTTTGGTTAAGATGTGACTGCCATCATCTTCGTGGTGGCCAGCAGCAGCTACAACATGGTCATCCGGG**atggtgttcaccctggaggacttcgtgggcgactgggagcagaccgccgcctacaacctggaccaggtgctggagcagggcggcgtgagcagcctgctgcagaacctggccgtgagcgtgacccccatccagcgcatcgtgcgcagcggcgagaacgccctgaagatcgacatccacgtgatcatcccctacgagggcctgagcgccgaccagatggcccagatcgaggaggtgttcaaggtggtgtaccccgtggacgaccaccacttcaaggtgatcctgccctacggcaccctggtgatcgacggcgtgacccccaacatgctgaactacttcggccgcccctacgagggcatcgccgtgttcgacggcaagaagatcaccgtgaccggcaccctgtggaacggcaacaagatcatcgacgagcgcctgatcacccccgacggcagcatgctgttccgcgtgaccatcaacagcgggagctccggtggtggcgggagcggaggtggaggctcgagtatgattgagaaacagctgcagaaggacaaacaggtctatagggcaacccataggctgctgctccttggcgccgataattccgggaaatccacgattgttaagcagatgaggatcctccatggagggtcaggcggaagtggaggcactagcggtatctttgaaacaaagtttcaggtggataaagtgaacttccacatgttcgatgttggcgggcaacgagatgaaagacggaagtggattcagtgctttaacgatgtgactgccatcatcttcgttgtggactctagcgactacaaccggctgcaagaagcacttaacgacttcaaatctatctggaataatcgatggctgagaaccataagcgtcatactgttcctgaataagcaggaccttctggctgaaaaggtattggctgggaagtccaagatagaggactacttccccgagtttgcccgctataccacacctgaggacgctaccccagagcctggtgaggatccacgtgtaacacgagctaaatacttcattcgcgacgaatttctccgcatctccactgcttctggggatggtcgtcactactgctatccccattttacctgtgccgtggacacggagaatgccaggagaattttcaacgactgtcgggatatcatccagcgcatgcatttgcggcaatacgagctgttggtttcaggaagcgga**gctactaacttcagcctgctgaagcaggctggagacgtggaggagaaccctggacctggatcgtttcgcatgattgaacaagatggattgcacgcaggttctccggccgcttgggtggagaggctattcggctatgactgggcacaacagacaatcggctgctctgatgccgccgtgttccggctgtcagcgcaggggcgcccggttctttttgtcaagaccgacctgtccggtgccctgaatgaactgcaggacgaggcagcgcggctatcgtggctggccacgacgggcgttccttgcgcagctgtgctcgacgttgtcactgaagcgggaagggactggctgctattgggcgaagtgccggggcaggatctcctgtcatctcaccttgctcctgccgagaaagtatccatcatggctgatgcaatgcggcggctgcatacgcttgatccggctacctgcccattcgaccaccaagcgaaacatcgcatcgagcgagcacgtactcggatggaagccggtcttgtcgatcaggatgatctggacgaagagcatcaggggctcgcgccagccgaactgttcgccaggctcaaggcgcgcatgcccgacggcgaggatctcgtcgtgacccatggcgatgcctgcttgccgaatatcatggtggaaaatggccgcttttctggattcatcgactgtggccggctgggtgtggccgaccgctatcaggacatagcgttggctacccgtgatattgctgaagagcttggcggcgaatgggctgaccgcttcctcgtgctttacggtatcgccgctcccgattcgcagcgcatcgccttctatcgccttcttgacgagttcttctgattcgaacatcAGGACAACCAGACCAACCAACCAGACCAACCGCCTGCAGGAGGCTCTGAACCTCTTCAAGAGCATCTGGAACAACAGGTTTGTGGAGTGACCGCCCACCCCCTGCGCTTGCCCAGGAGGCCCTGGTCTGCACTGTTTATAGAGAAGAACCCCGTGCAAGCATTCCAGACCCCTGGCCGAAAGCGCGCTTCTCCCAAGCATTCACACGGCCTCCCTTCTTGTAGATGGCTGCGCACCATCTCTGTGATCCTGTTCCTCAACAAGCAAGATCTGCTCGCTGAGAAAGTCCTTGCTGGGAAATCGAAGATTGAGGACTACTTTCCAGAATTTGCTCGCTACACTACTCCTGAGGATGGTGTGTATGGCTTCCACTCTTGCTGGCTGTTCATTGCGGTGGTTCTTTTTCAAACGGTCAGGCTGAAAACCCCCATCCCCCTCCCACCACCAAACCATAAAGGATCTATAAGAGAAGCAAGAAAAACGCACTCCCACTAATTCTCATATGGAAAAATCAGGGTTTTGAAGACTTCAGGAGCTACAGAGATGCTAGCACCCCAGCTCTGCTTGAATTTTAAATTACATTAATATGTATTCCCTTTTTATATAGCTACTCCCGAGCCCGGAGAGGACCCACGCGTGACCCGGGCCAAGTACTTCATTCGAGATGAGTTTCTGGTGAGTCGAGCCTGTCTTTAGTTTCCTCTCTTGTTCCTCCTCTTTTTCTCATGGATGTAAATTTACTTAATTCCAAATTCAGGGGTTCAGCTACCCAGTT |

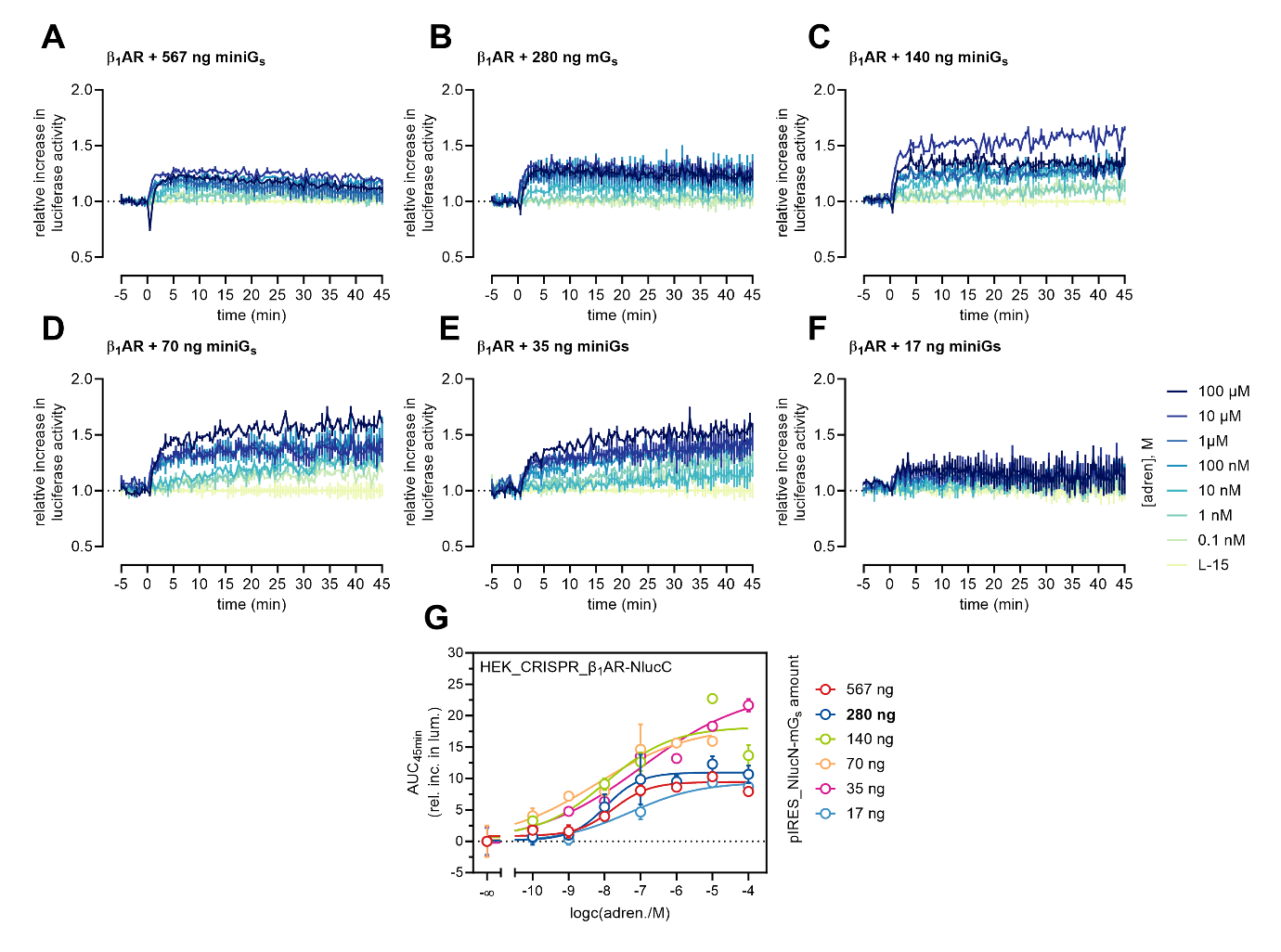

**Figure S2** – Optimisation of HEK_CRISPR_ β_1_AR-NlucC cells with overexpressed NlucN-mG_s_. (**A-F**) Kinetic response of varied isoprenaline concentrations with the split-luciferase assay using (A) 567 µg, (B) 280 ng, (C) 140 ng, (D) 70 ng, (E) 35 ng or (F) 17 ng NlucN-mG_s_ plasmid. Baseline L-15 = buffer (0% response). (**G**) Concentration response curves of adrenaline in transiently transfected HEK_CRISPR_ β_1_AR-NlucC cells with indicated amounts of pIRES_puro_3_NlucN-mG_s_ plasmid DNA. In further experiments, 280 ng of mG_s_ plasmid DNA was used for transfection. Data represent the mean ± SEM of *n* = 1 experiment carried out in triplicate.

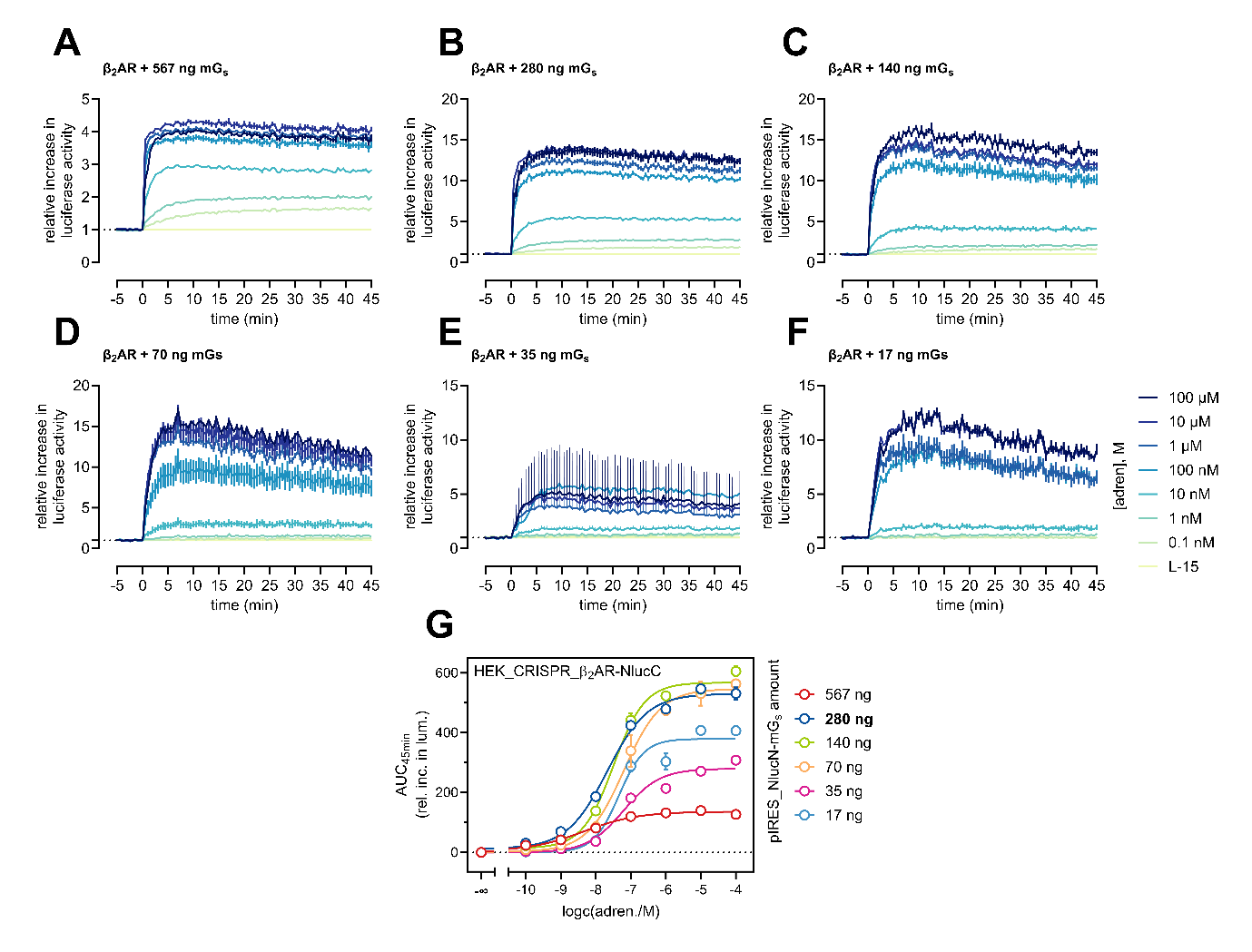
**Figure S3** – Optimisation of HEK_CRISPR_ β_2_AR-NlucC cells with overexpressed NlucN-mG_s_. (**A-F**) Kinetic response of varied isoprenaline concentrations with the split-luciferase assay using (A) 567 µg, (B) 280 ng, (C) 140 ng, (D) 70 ng, (E) 35 ng or (F) 17 ng NlucN-mG_s_ plasmid. Baseline L-15 = buffer (0% response). (**G**) Concentration response curves of adrenaline in transiently transfected HEK_CRISPR_ β_2_AR-NlucC cells with indicated amounts of pIRES_puro_3_NlucN-mG_s_ plasmid DNA. In further experiments, 280 ng of mG_s_ plasmid DNA was used for transfection. Data represent the mean ± SEM of *n* = 1 experiment carried out in triplicate.

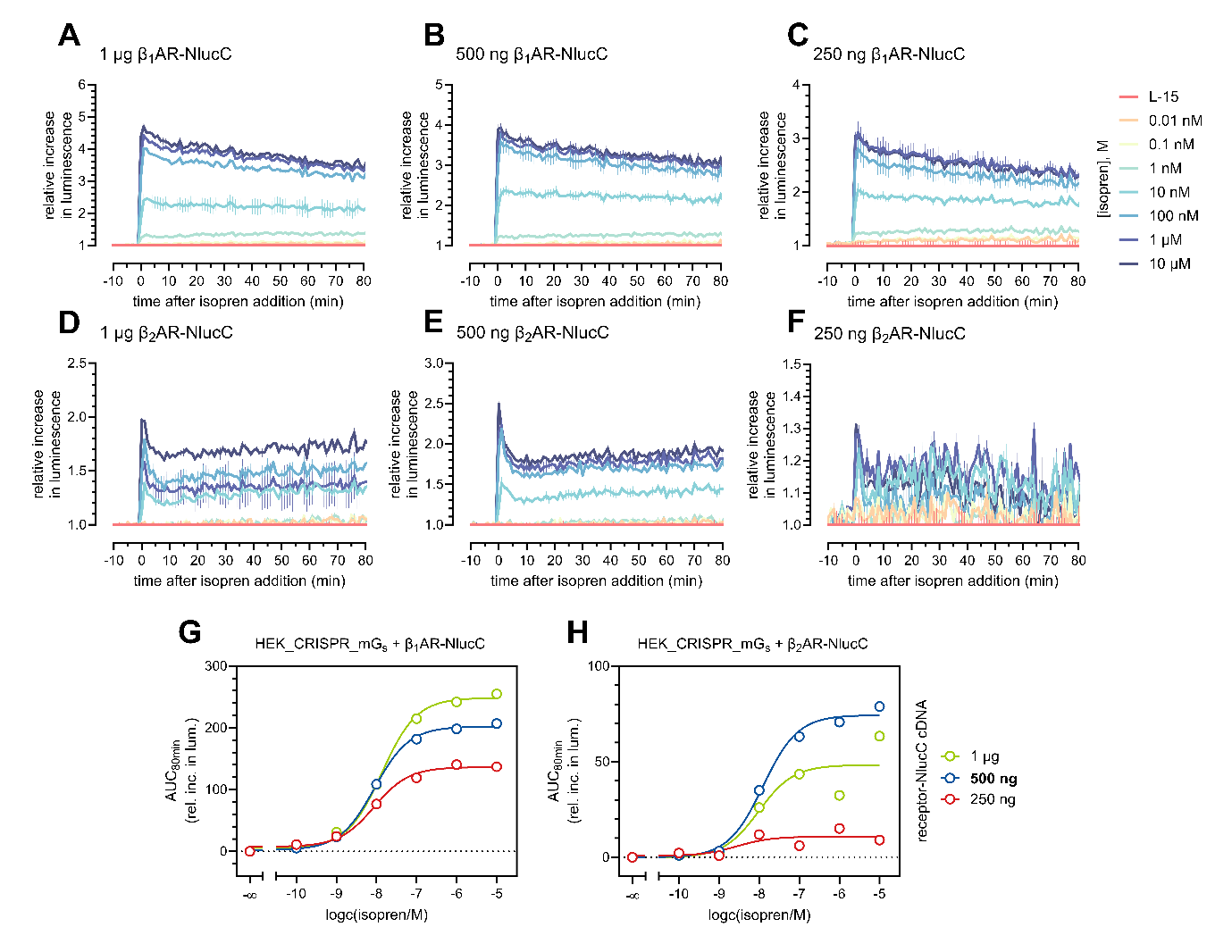

**Figure S4** – Optimisation of HEK_CRISPR_NlucN-mG_s_ cells with overexpressed β_1,2_AR-NlucC. (**A-F**) Kinetic response of varied isoprenaline concentrations with the split-luciferase assay using (A,D) 1 µg, (B, E) 500 ng or (C, F) 250 ng receptor-NlucC plasmid using 5 µM coelenterazine h. Baseline L-15 = buffer (0% response). (**G, H**) Concentration response curves of isoprenaline response in transiently transfected HEK_CRISPR_NlucN-mG_s_ cells with indicated amounts of p3.1_β_1,2_AR-NlucC plasmid DNA. In further experiments, 500 ng of receptor plasmid DNA was used for transfection. Data represent the mean ± SEM of *n* = 1 experiment carried out in triplicate.

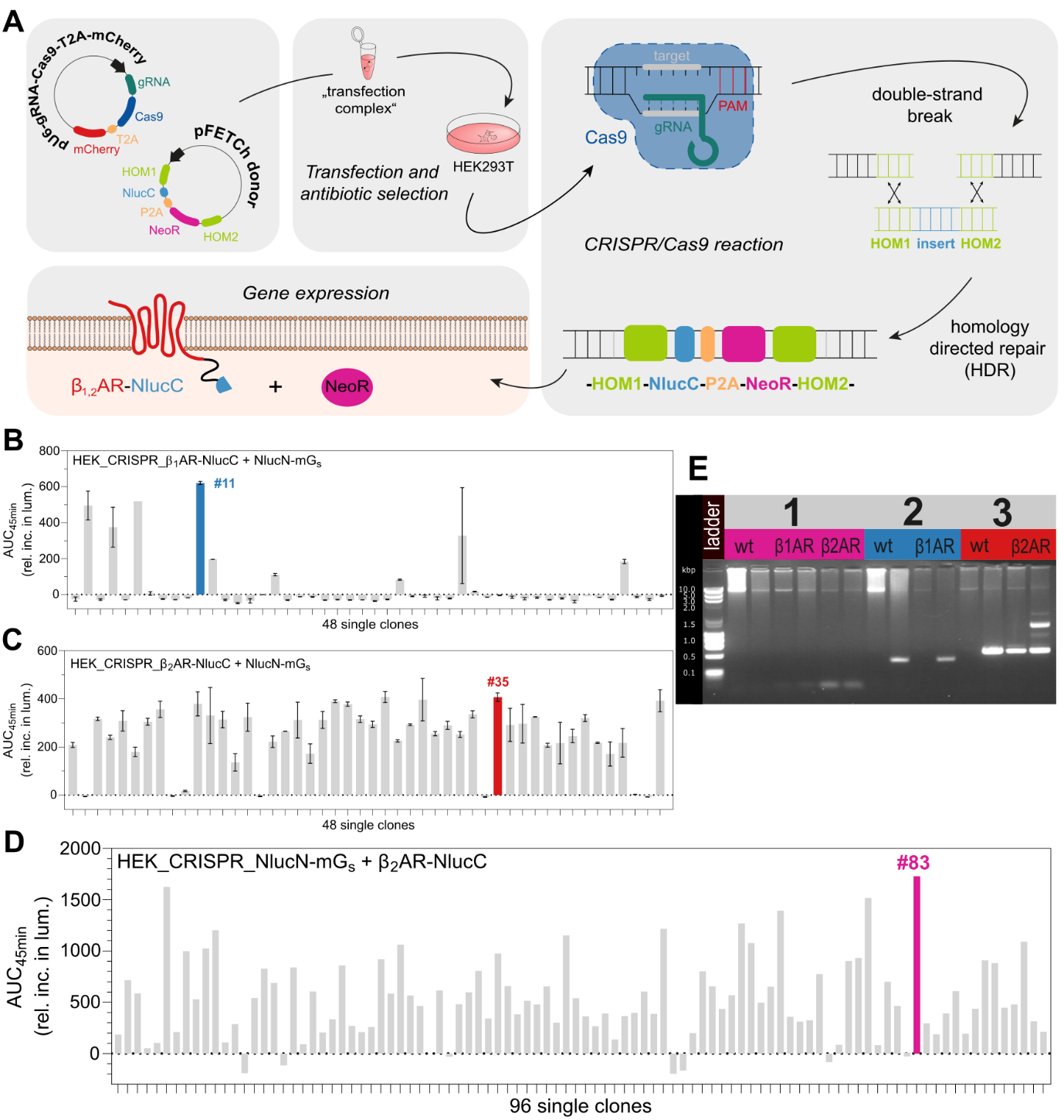

**Figure S5** – Schematic representation of the generation and verification of HEK293T cells expressing β_1,2_AR-NlucC or NlucN-mG_s_ fusion proteins under endogenous promotion. **A**) Workflow of the implemented CRISPR/Cas9 technique. Single clones were obtained from genome-edited HEK293T cells and transiently transfected with overexpressed split luciferase partners. The clones were screened in response to 10 μM adrenaline of **B**) β_1_AR-NlucC, **C**) β_2_AR-NlucC, both with NlucN-mG_s_ in duplicate, or **D**) NlucN-mG_s_ with β_2_AR-NlucC (*n* = 1). β_1_AR-NlucC clone 11, β_2_AR-NlucC clone 35 and NlucN-mG_s_ clone 83 were selected for further experiments. **E**) PCR products from extracted genomic DNA from wildtype HEK293T (wt), CRISPR/Cas9 tagged β_1_AR-NlucC (β1AR) or β_2_AR-NlucC (β2AR). Primers were varied to extract NlucC (1; 70 bp product), β_1_AR-NlucC (2; untagged 600 bp, tagged 1,600 bp) or β_2_AR-NlucC (3 untagged 700 bp, tagged 1,700 bp).

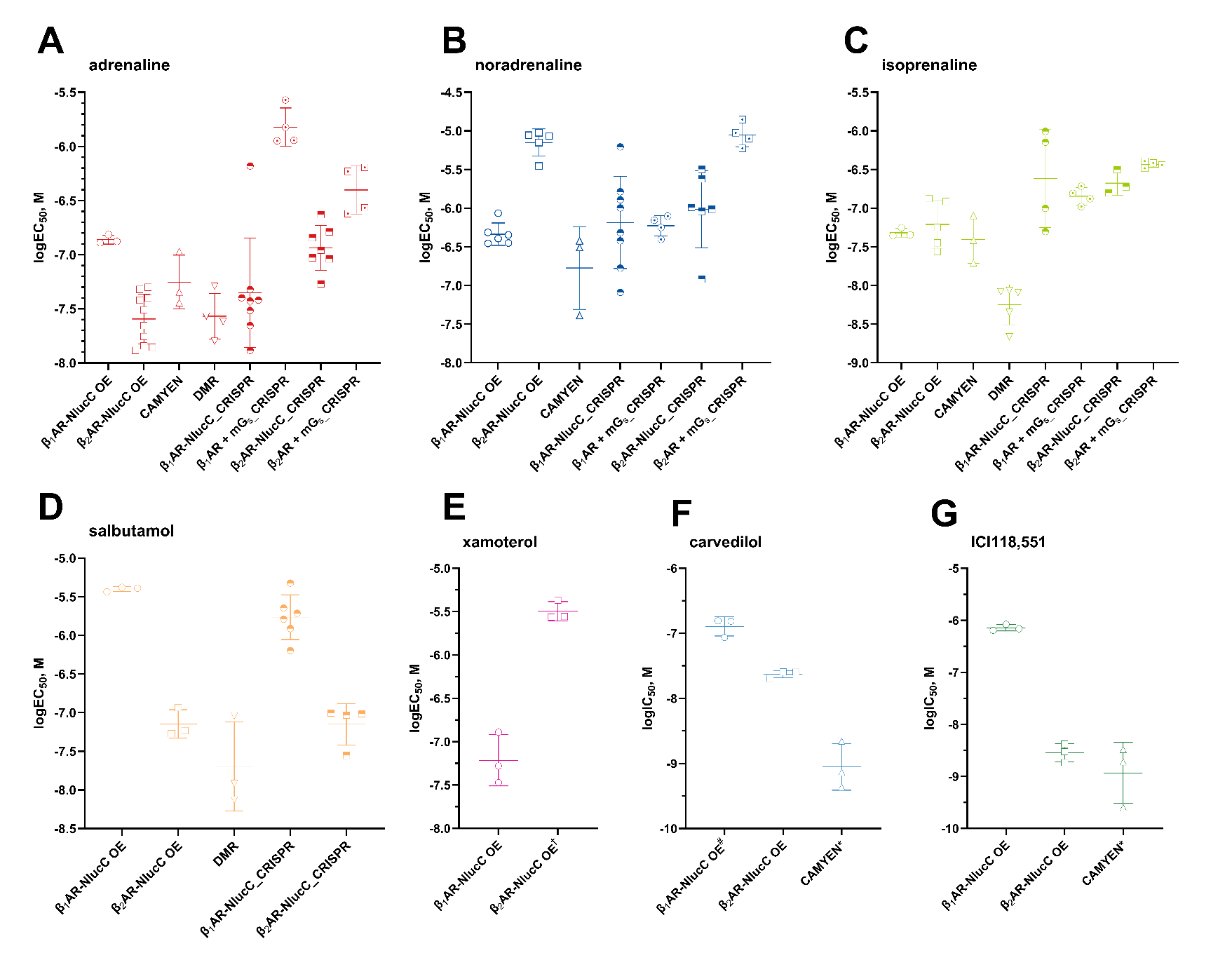

**Figure S6** – Potency values for individual experiments. Different ligands are shown with different experimental systems for (**A**) adrenaline, (**B**) noradrenaline, (**C**) isoprenaline, (**D**) salbutamol, (**E**) xamoterol, (**F**) carvedilol and (**G**) ICI118,551. IC_50_ values were determined in the presence of 100 nM adrenaline. OE = overexpressed. ^†^measured as displacement of 100 nM adrenaline, ^#^measured as logEC_50_ without agonist, *measured in the presence of 100 nM isoprenaline. Data are shown as mean ± standard deviation and *n* ≥ 3 individual experiments performed in triplicate.

**Table S4 –** Potencies (pEC_50_), efficacies (E_max_) and Δlog(E_max_/EC_50_) relative to adrenaline obtained by mG_s_ recruitment. Either HEK293T cells that stably expressed NlucN-mG_s_ and β_1,2_AR-NlucC fusion proteins (Overexpressed), genome edited HEK293T_CRISPR_β_1,2_AR-NlucC cells transiently expressing the NlucN-mG_s_ fusion protein (CRISPR_receptor) or genome edited HEK293T_CRISPR_ NlucN-mG_s_ cells transiently expressing the β_1,2_AR-NlucC fusion proteins (CRISPR_mG_s_) were used. Agonist responses were normalized to L-15 as solvent control (0%) and to the maximal response elicited by 10 µM adren, or 100 µM adren for overexpressed β_1_AR. Data represent means ± SEM from at least three independent experiments (n ≥ 3), performed in triplicate, *data were obtained from a three-parameter fit, and n.a. = not applicable, where the ligand did not yield a full concentration-response curve up to 100 µM.

|  | **Overexpressed** | | | **CRISPR_receptor** | | | **CRISPR_mG_s_** | | |
| --- | --- | --- | --- | --- | --- | --- | --- | --- | --- |
| Cpd. | pEC_50_ | E_max_ | Δlog($\frac{E_{max}}{{EC}_{50}}$) | pEC_50_ | E_max_ | Δlog($\frac{E_{max}}{{EC}_{50}}$) | pEC_50_ | E_max_ | Δlog($\frac{E_{max}}{{EC}_{50}}$) |
| **β_1_AR** | | | | | | | | | |
| adren | 6.86 ± 0.02 | 100 | 0 | 6.60 ± 0.12 | 100 | 0 | 5.82 ± 0.08 | 100 | 0 |
| noradren | 6.21 ± 0.06 | 115 ± 17 | 0.69 ± 0.10 | 6.18 ± 0.20 | 109 ± 7 | 0.44 ± 0.23 | 6.23 ± 0.06 | 99 ± 8 | -0.41 ± 0.11 |
| isopren | 7.32 ± 0.03 | 110 ± 1 | -0.42 ± 0.03 | 6.61 ± 0.28 | 107 ± 9 | 0.01 ± 0.29 | 6.85 ± 0.05 | 114 ± 11 | -0.98 ± 0.12 |
| salbut | 5.4 ± 0.02 | 37 ± 1 | 1.03 ± 0.03 | 5.76 ± 0.11 | 37 ± 6 | 0.37 ± 0.14 | n.a. | n.a. | n.a |
| **β_2_AR** | | | | | | | | | |
| adren | 7.55 ± 0.06 | 100 | 0 | 6.94 ± 0.07 | 100 | 0 | 6.40 ± 0.10 | 100 | 0 |
| noradren | 5.12 ± 0.05 | 85 ± 2 | 2.36 ± 0.08 | 6.01 ± 0.19 | 61 ± 7 | 0.69 ± 0.21 | 5.05 ± 0.07 | 51 ± 4 | 1.05 ± 0.12 |
| isopren | 7.21 ± 0.12 | 107 ± 5 | 0.37 ± 0.15 | 6.68 ± 0.07 | 112 ± 12 | 0.30 ± 0.14 | 6.43 ± 0.02 | 124 ± 3 | 0.06 ± 0.10 |
| salbut | 7.15 ± 0.09 | 82 ± 1 | 0.32 ± 0.11 | 7.15 ± 0.12 | 67 ± 6 | -0.39 ± 0.15 | 6.75 ± 0.15* | 23 ± 4* | -1.03 ± 0.21* |
